# Root gene regulatory network analysis reveals roles for defense hormones and secondary metabolites in establishment of disease-suppressive soil

**DOI:** 10.64898/2026.10.22.760950

**Authors:** Run Qi, Ivona Glavincheska, Jelle Spooren, Xiaoyan Liu, Marit Van Laatum, Francel Verstappen, Iris F. Kappers, Roeland L. Berendsen, Corné M.J. Pieterse, Saskia C.M. Van Wees

## Abstract

- Plants recruit beneficial microbiota in their rhizosphere to alleviate stresses. In *Arabidopsis thaliana*, foliar infection by *Hyaloperonospora arabidopsidis* (*Hpa*) has been shown to establish a disease-suppressive microbiome that promotes protection against *Hpa* infection of the subsequent plant generation grown in this conditioned soil - a phenomenon known as soil-borne legacy (SBL). The root responses involved in SBL establishment have remained largely unclear.
- In this study, time-series RNA-seq of roots from *Hpa*-infected plants grown in natural soil was conducted over seven days post-inoculation. A gene regulatory network of the responding roots was constructed. The role of predicted key components was verified in follow-up assays.
- Root transcriptional reprogramming started within one day after foliar infection. Genes associated with salicylic acid (SA)-mediated defense were upregulated, while those linked to secondary metabolism and jasmonic acid (JA) responsiveness were downregulated. SA accumulation increased at 1 dpi, and JA decreased by 3 dpi. Transcriptional regulation occurred primarily in the tips of the roots. Mutant analyses confirmed that SA, JA, indole glucosinolates, camalexin, and flavonoids contribute to SBL establishment.
- Our findings reveal rapid systemic transcriptional reprogramming in roots upon foliar pathogen attack, uncovering novel roles of hormone signaling and secondary metabolism in shaping a disease-suppressive soil beyond their known defense functions.

## Introduction

Soil harbors billions of microbes, among which plants selectively promote beneficial ones to support their growth and health. Under stress conditions, plant-derived cry-for-help signals shape specific microbial consortia that can persist in the soil, establishing a protective legacy for future plant generations. This phenomenon, known as soil-borne legacy (SBL) (Bakker *et al*. 2018), plays a crucial role in plant resilience. When *Arabidopsis thaliana* (hereafter Arabidopsis) leaves are infected by the downy mildew pathogen *Hyaloperonospora arabidopsidis* (*Hpa*), the rhizosphere and phyllosphere microbiomes undergo distinct changes that underly a disease-suppressive SBL. Spooren *et al*. (2026) demonstrated that the disease suppressiveness of the SBL soil is caused by living microbes. Specific members of the enriched microbiota have been shown to enhance plant resistance against *Hpa* (Berendsen *et al*. 2018; Goossens *et al*. 2023), likely by priming the plant’s immune system (Vismans *et al*. 2022).

*Hpa* is an obligate biotrophic pathogen that relies on living Arabidopsis host cells to complete its life cycle (Coates and Beynon 2010). The plant hormone salicylic acid (SA) is crucial in coordinating induction of defense responses that mitigate *Hpa* infection. SA biosynthesis mutant *sid2* and SA signaling mutant *npr1* exhibit enhanced disease susceptibility compared to wild type, whereas SA-overaccumulating mutants *dmr6* and *nac61nac90* show reduced *Hpa* infection (Delaney *et al*. 1995; Nawrath and Métraux 1999; Van Damme *et al*. 2008; Hickman *et al*. 2019). Other hormones, such as abscisic acid (ABA) and cytokinin (CK), also contribute to *Hpa* resistance by promoting callose deposition and enhancing SA-dependent defense responses, respectively (Ton *et al*. 2005; Argueso *et al*. 2012). SA is essential for systemic acquired resistance, which enhances defense in distant tissues and provides broad-spectrum protection against (hemi-)biotrophic pathogens like *Hpa* and *Pseudomonas syringae* (Métraux *et al*. 1990; Delaney *et al*. 1994). While SA signaling has been well-studied in leaves, its role in roots remains underexplored. Although SA has been implicated in rhizosphere microbiome assembly (Lebeis *et al*. 2015), the regulatory mechanisms of the SA signaling in root tissues are still largely unknown.

In contrast to SA, jasmonic acid (JA) suppresses defense against *Hpa*, as evidenced by the enhanced resistance of the JA-insensitive mutant *coi1* and by *Hpa* effectors HaRxL44 and HaRxL10, which activate JA signaling to antagonize SA-mediated defense and facilitate infection (Caillaud *et al*. 2013; Murmu *et al*. 2014; Anderson *et al*. 2019). However, JA is critical for defense against necrotrophic pathogens and insect herbivores (Pieterse *et al*. 2012) and plays a key role in induced systemic resistance (ISR) triggered by beneficial rhizosphere microbes or endophytic root colonizers (Pieterse *et al*. 2014; 2021). This form of ISR primes the entire plant for enhanced JA-dependent defenses, providing broad-spectrum resistance in roots and shoots. Notably, ISR induced by *Pseudomonas simiae* WCS417 is also effective against *Hpa*, through priming for callose deposition, which forms a physical barrier to infection (Ton *et al*. 2002; Van der Ent *et al*. 2008). JA signaling also influences rhizosphere microbiome composition, as evidenced by microbial community shifts in JA signaling mutants or after exogenous foliar application of methyl jasmonate (MeJA), which alters root exudation profiles (Carvalhais *et al*. 2013; Carvalhais *et al*. 2015; Carvalhais *et al*. 2017).

In response to pathogen or insect attack, Arabidopsis increases the production and/or release of antimicrobial secondary metabolites, like phenylpropanoids, and S-containing phytoalexins and glucosinolates. Phenylpropanoids are produced via a complex metabolic pathway that generates a wide variety of different compounds, including lignin, flavonoids, and coumarins, which share common biosynthetic precursors (Vogt 2010). Lignin reinforces plant cell walls, acting as a hydrophobic barrier to pathogen invasion (Lee *et al*. 2019). Flavonoids scavenge reactive oxygen species (ROS) under stress and can also induce microbial gene expression, such as the *nod* genes in rhizobia, facilitating symbiosis (Djordjevic *et al*. 1987). Flavonoid overaccumulation has been shown to alter rhizosphere microbiota composition in Arabidopsis and buckwheat (He *et al*. 2022; Wu *et al*. 2025). Another group of phenylpropanoids, coumarins, are extensively studied for their roles in iron acquisition and antimicrobial defense. Coumarins are exuded in response to iron deficiency to mobilize iron for uptake by the roots. Beneficial rhizobacteria such as *Pseudomonas simiae* WCS417 and *Pseudomonas capeferrum* WCS358 tolerate the antimicrobial coumarin scopoletin, whereas soil-borne pathogens are suppressed (Stringlis *et al*. 2018). The coumarin biosynthesis genes *MYB72* and *F6’H1* are essential for *Hpa*-induced SBL formation, as *myb72* and *f6’h1* mutants fail to establish a protective SBL, coinciding with altered root exudation and microbiome composition (Vismans *et al*. 2022). These findings suggest that coumarins play a critical role in shaping the root microbiome, although it is not known whether these coumarin-dependent changes in the microbiome are causally related to SBL establishment.

The S-containing compound camalexin is the main phytoalexin of Arabidopsis, and is crucial for resistance against necrotrophic fungi such as *Botrytis cinerea* and *Alternaria brassicicola* (Thomma *et al*. 1999; Ferrari *et al*. 2003; Van Wees *et al*. 2003). Although camalexin is also induced upon *Hpa* infection, it plays a minimal role in defense against *Hpa* (Mert-Türk *et al*. 2003). Camalexin can also influence the rhizosphere microbiome composition. Mutations in *CYP71A* genes, which impair camalexin biosynthesis, reduce rhizosphere sulfatase activity and diminish the plant growth-promoting effects of *Pseudomonas fluorescens* CH267 (Koprivova *et al*. 2019). Moreover, it can reduce the swimming motility of rhizosphere colonizing bacteria, resulting in the reduction of root colonization (Zhou *et al*. 2025).

Glucosinolates, another major group of S-containing secondary metabolites, become biologically active upon cell damage, breaking down into antimicrobial and insecticidal isothiocyanates. There are two main types of glucosinolates: indole glucosinolates (IGS), derived from tryptophan, and aliphatic glucosinolates (AGS), derived from methionine. Their breakdown products inhibit plant pathogens such as *Erwinia carotovora*, *Verticillium longisporum,* and *Sclerotinia sclerotiorum* (Brader *et al*. 2001; Tierens *et al*. 2001; Witzel *et al*. 2013). IGS can also promote callose deposition (Clay *et al*. 2009) and limit colonization of the roots by non-pathogenic *Trichoderma* strains (Poveda 2021).

Several key transcription factors (TFs) regulate secondary metabolite biosynthesis in Arabidopsis. The MYC TFs activate JA-responsive genes, while JAZ proteins act as repressors that are degraded upon JA signaling activation, thereby acting as molecular switches. ABA contributes to secondary metabolism partly by enhancing the MYC branch of the JA pathway, which is regulated by MYC2 and activated upon herbivory (Vos *et al*. 2013). Insect feeding on the triple MYC mutant *myc2/3/4* fails to induce IGS production, whereas *jazQ* and *jazD* mutants hyperaccumulate these secondary metabolites (Schweizer *et al*. 2013; Guo *et al*. 2022). The induction of flavonoid biosynthesis genes upon wounding is also dependent on MYC3 (Wang *et al*. 2021a). MYB TFs additionally regulate specific branches of secondary metabolism. MYB34/51/122 are essential for IGS biosynthesis (Frerigmann and Gigolashvili 2014), while MYB28, MYB29, and MYB76 regulate AGS biosynthesis (Gigolashvili *et al*. 2007; 2008). Other MYB TFs modulate phenylpropanoid metabolism, influencing suberin, lignin, flavonoid, or coumarin biosynthesis (Liu *et al*. 2015a; Stringlis *et al*. 2018). While SA plays a relatively minor role in secondary metabolism, it can induce biosynthesis of camalexin and the IGS 4MO-I3M in Arabidopsis under specific conditions (Mikkelsen *et al*. 2003; Frerigmann and Gigolashvili 2014). TFs such as WRKY33 and MYB51, which regulate the biosynthesis of 4MO-I3M and camalexin biosynthesis, are themselves influenced by SA, JA and/or ABA signaling (Frerigmann and Gigolashvili 2014; Liu *et al*. 2015b; Tao *et al*. 2022).

Our research aimed to identify root responses triggered by foliar *Hpa* infection that contribute to SBL establishment. To this end, we monitored root transcriptional changes over a seven-day time course following leaf infection. While previous studies have explored shifts in rhizosphere microbiome composition following interactions with above-ground pathogens (Berendsen *et al*. 2018; Yuan *et al*. 2018), herbivores (Friman *et al*. 2021; Wang *et al*. 2021b; Yang *et al*. 2025), or abiotic stresses such as reduced light or drought (Hou *et al*. 2021; Munoz-Ucros *et al*. 2022; Fang *et al*. 2025), little is known about the systemically induced transcriptional changes in roots. With the advancement of next-generation sequencing, gene regulatory network (GRN) analysis has become a powerful tool to decipher complex gene expression programs and has been successfully applied to study plant development and immune responses (Windram and Denby 2015; Alvarez *et al*. 2021). Here, we constructed a GRN model of Arabidopsis root transcriptional reprogramming associated with the establishment of disease-suppressive soil following foliar *Hpa* infection. Validation experiments based on GRN-derived predictions highlighted the roles of SA, JA, and the secondary metabolites 4MO-I3M, camalexin, and flavonoids in establishing a disease-suppressive soil.

## Materials and Methods

### Plant material and growth conditions

All wild-type, mutant, and transgenic *Arabidopsis thaliana* plants used in this study were in the Columbia accession (Col-0) background. The following T-DNA insertion mutants and transgenic lines were obtained from the Nottingham Arabidopsis Stock Centre: *wrky33-1* (SALK_006603) (Zheng *et al*. 2006), *cyp81f2* (SALK_073776) (Pfalz *et al*. 2009), *cyp81f4* (SALK_024438) (Pfalz *et al*. 2011), *tt3-5*, and *tt4-3* (Peer *et al*. 2001). Mutant lines *pad3-1* (Zhou *et al*. 1999), *myb28/29* (*myb28-1 myb29-1)* double mutant (Sønderby *et al*. 2007), *npr1-1* (Cao *et al*. 1997), *sid2-1* (Wildermuth *et al*. 2001), and *myc2* (*jin1-7*) (Berger *et al*. 1996) were already present in our laboratory. GUS reporter lines *WRKY33pro:GUS* (Lippok *et al*. 2007) and *JAZ10pro:GUS* (Acosta *et al*. 2013) were kindly provided by Paul Schulze-Lefert and Edward Farmer, respectively. Arabidopsis lines used in this study are also summarized in Table **S1**. Plants were grown as previously described in detail (Vismans *et al*. 2022). In brief, seeds were placed into 0.2% water agar and stratified at 4°C for 48 h prior to sowing on natural field soil, collected from the Reijerscamp nature reserve in the Netherlands where endemic Arabidopsis grows. The key physicochemical properties of this soil have been previously described in Berendsen *et al*. (2018). To ensure spatial consistency and reduce algal growth, the soil surface was covered with a circular cutout of a plastic micropipette-tip holder (Greiner Bio-One, 0.5– 10 µl, item no. 771280), which served as a reusable sowing template (Vismans *et al*. 2021). Approximately 40 seeds were sown by pipetting them into each pot. The pots were then placed in a climate-controlled plant growth chamber under a 10-h day (100 μmol m^−2^ s^−1^) and 14-h night cycle at 21°C, 70% relative humidity.

### *In vitro* plate assay with *Hpa* or MeJA

Arabidopsis seeds were surface-sterilized using the gas sterilization method described by Van Wees *et al*. (2013), then sown on square plates containing MS agar-solidified medium (with vitamins, no sucrose, pH 5.8). Plates were placed at 4°C in darkness for 2 days to synchronize germination. After stratification, plates were positioned vertically and incubated in the same growth condition as described above. After two weeks of growth, *Hpa* spores were inoculated onto cotyledons or true leaves using an electronic pipette, applying three droplets of 0.3 μl of 50 spores/ul per leaf, following the method described by Goossens *et al*. (2023). Plates were sealed with Parafilm (Sigma) and returned to the same growth conditions for seven days to allow disease development. For MeJA treatment, seedlings were transferred to 6-well plates filled with 50 μM MeJA solution and submerged for 3 h.

### The “tea bag system” to grow intact and clean roots in soil

A modified pot setup, referred to as the "tea bag system," was developed to cultivate intact and soil-free roots for studying root-microbe interactions in a soil environment (Fig. **S1a,b**). In this system, standard 60-ml pots were cut in half and each side was sealed with SEFAR® NITEX 03-100/44 nylon mesh (100-µm pore opening). When reassembled, this created a mesh-enclosed, pocket-like space. Arabidopsis seeds were germinated and grown within this enclosed, non-soil zone between the nylon layers, enabling root development without direct contact with soil. Meanwhile, microbial exposure of the roots was maintained through microbes migrating from the surrounding soil in the rest of the pot.

### Experimental setup of induction of a disease-suppressive SBL

*Hpa* isolate Noco2 was used for SBL induction and disease assays. To cultivate the *Hpa* inoculum, 10-day-old Arabidopsis Col-0 seedlings were spray-inoculated with a spore suspension of *Hpa* Noco2 (50 spores/µl in tap water) using an airbrush. Plants were maintained under high humidity, and spores were harvested 7 days post inoculation (dpi) by washing off sporulating leaves through shaking of the harvested shoots for 15 s in tap water using a vortex. The resulting spore suspension was used to maintain the *Hpa* culture by inoculating a new batch of seedlings, or for experiments to induce a disease-suppressive SBL, following the protocol previously described (Vismans *et al*. 2022). In brief, two weeks after sowing (as described above), half of the pots were sprayed with a 50 spores/µl suspension of *Hpa* Noco2 until small droplets formed on the leaves (approximately 0.5 ml per pot). The remaining pots were sprayed with tap water as a mock treatment. These plants, referred to as the “conditioning phase”, were either used directly for various downstream experiments (e.g., root RNA-seq), or left in place for one week to condition the soil for subsequent SBL testing. For SBL experiments, the aboveground plant parts were removed using a razor blade, and a new population of seeds was sown on the conditioned soil. Two weeks later, these “response phase” plants were inoculated with *Hpa* using the same method as above. Disease severity was quantified by counting spores and normalizing to shoot fresh weight, as described by Goossens *et al*. (2023). Briefly, shoots were harvested, weighed, and placed in 3-6 ml of water, vortexed for 15 s, and spores were counted in three 2-μl droplets under a microscope. The average spore count per plant was used for analysis. For SA dipping treatment during the conditioning phase, the procedure was identical to the *Hpa* treatment, except that shoots were dipped in a 1 mM SA solution containing 0.015% CoatOSil 77 (Momentive).

### Experimental setup and RNA isolation for RNA-seq

Col-0 plants were cultivated and treated with *Hpa* or water (mock) as described above. For the time-series RNA-seq experiment, roots from the "conditioning phase" were harvested from individual *Hpa*- or mock-treated pots at each of the following time points post-treatment: 1, 2, 3, 4, 5, 6, and 7 d. Each sample consisted of roots from approximately thirty 2- to 3-week-old plants growing in a single pot. Roots were collected in a sieve, rinsed under running water for approximately 20 s, blotted dry with tissue paper, flash-frozen in liquid nitrogen, and finely ground using a TissueLyser II (Qiagen). RNA was extracted following the protocol described by Hickman *et al*. (2017), using the RNeasy Plant Mini Kit (Qiagen) with an included DNase treatment step according to the manufacturer’s instructions. RNA integrity (RIN) was assessed using an Agilent Plant RNA Nano Chip, and only samples with RIN > 7 were selected for cDNA library preparation and sequencing. In total, 4 to 6 biological replicates per treatment and time point were independently processed and sequenced. For 1 dpi, mock *n* = 5 and *Hpa n* = 6; at 2 dpi, mock *n* = 4 and *Hpa n* = 6; at 3 dpi, mock *n* = 5 and *Hpa n* = 6; at 4 dpi, mock *n* = 4 and *Hpa n* = 6; at 5 dpi, mock *n* = 6 and *Hpa n* = 6; at 6 dpi, mock *n* = 4 and *Hpa n* = 6; and at 7 dpi, mock *n* = 5 and *Hpa n* = 6. Library preparation and sequencing were performed at the Oklahoma Medical Research Foundation NGS Core using an Illumina NovaSeq S4 platform, generating 20 million 150 bp paired-end reads per sample.

### RNA-seq analysis

Sequencing reads were aligned to the Arabidopsis genome (TAIR version 10) using Hisat2. Read counts were summarized over annotated gene models using featureCounts, based on the Arabidopsis annotation file provided by Ensembl Plants (Anders *et al*. 2015), and normalized using the DESeq2 R package (Love *et al*. 2014). Genes differentially expressed between mock and *Hpa* treatment at individual timepoints (false discovery rate < 0.05) were identified using DESeq2 with default settings (Kim *et al*. 2019). To identify genes whose transcript levels differed significantly over time between the mock and *Hpa* treatments, we applied the approach previously described (Hickman *et al*. 2017). Briefly, for each gene, a negative binomial generalized linear model (GLM) was fit to the normalized gene counts with treatment and time as covariates. We then used ANOVA to compare the full model to a reduced model without the treatment variable. To adjust for multiple comparisons, we applied the Bonferroni correction to the resulting *P* values. Genes with an adjusted *P* value < 0.05 at one or more timepoints were classified as differentially expressed genes (DEGs). For each DEG, the time point of first differential expression was predicted as described previously (Hickman *et al*. 2017).

### Clustering of gene expression profiles and GO term enrichment analysis

Clustering of DEGs was performed using SplineCluster (Heard *et al*. 2006) and DREM (Schulz *et al*. 2012). With Splinecluster, DEGs generated by RNA-seq analysis were clustered with a prior precision value of 10^-5^ with the variance stabilizing transformation (VST) method provided by DESeq2 (Love *et al*. 2014). An additional reallocation function, which redistributes cluster outliers into more appropriate clusters, was also performed. DREM was run using the same input data as SplineCluster and without building the network with TF interactions. Gene ontology (GO) term enrichment analysis was done using the enrichGO function in clusterProfiler v4.2.1 (Wu *et al*. 2021) with the following parameter changes. The OrgDb used was GO slim file provided by TAIR, ‘ont’ was set to ‘BP’, the pAdjustMethod was set to ‘BH’, and the pvalueCutoff were set to 0.01. The enrichment analysis was executed for each Splinecluster cluster or DREM path separately. To reduce redundancy among enriched GO terms, we applied the pairwise_termsim function from the enrichplot v1.24.4 in R (Yu 2021).

### Promoter motif and transcription factor binding site analyses

For promoter motif analysis, the promoter sequences, defined as the 1500 bp upstream of the predicted transcription start site, were retrieved from TAIR version 10. In total, promoter motif information for 568 Arabidopsis TFs was retrieved from the JASPAR database (Rauluseviciute *et al*. 2024). Simple enrichment analysis (SEA), developed by MEME Suite, was applied to the promoters of all genes present in a given Splinecluster cluster or DREM path, using the 568-motif information as reference, with all other parameters kept as default (Bailey and Grant 2021). For TF binding site analysis, essentially the method as described by Aerts *et al*. (2024) was followed. TF-gene interactions were inferred from DAP-seq (DNA affinity purification sequencing) experiments, which provide the genome-wide binding profiles of *in-vitro*-expressed TFs (O’Malley *et al*. 2016). DAP-seq peaks for 347 Arabidopsis TFs with a FRiP (fraction of reads in peaks) score ≥5% were retrieved from the Plant Cistrome DB. The enrichment of TF targets within co-expressed gene clusters or paths was assessed using the hypergeometric distribution.

### GUS assay

GUS staining was performed on 2-week-old seedlings grown in the above-described *in vitro* plate system and the tea bag system. Whole plants were submerged in GUS staining solution (1 mM X-Gluc, 100 mM NaPi buffer, pH 7.0, 10 mM EDTA, and 0.1% (v/v) Triton X-100), followed by vacuum infiltration and incubation at 37°C for 4 h. Chlorophyll was then removed with 96% ethanol (Spoel *et al*. 2003). Images were captured using a Zeiss microscope equipped with a camera.

### Hormone measurements

Plants were cultivated and treated with *Hpa* as described above. Roots and shoots from the "conditioning phase" were harvested from individual *Hpa*- or mock-treated pots at 0, 1, 3 and 7 days post treatment. Directly upon harvesting, root samples were flash frozen in liquid nitrogen and finely ground with a TissueLyser II (Qiagen). Ground samples (20±1 mg) were extracted with 1 ml of 10% methanol containing 100 nM stable isotope-labeled internal standards ([^2^H_6_]-(±)-JA, [^2^H_2_]-(−)-JA-Ile, [^2^H_4_]-SA). Authentic unlabeled standards ((−)-JA, (−)-JA-Ile, and SA) and stable isotope-labeled internal standards ([²H₆]-(±)-JA, [²H₂]-(−)-JA-Ile, and [²H₄]-SA) were provided by Olchemim Ltd. (Olomouc, Czech Republic).

Phytohormones were extracted from samples using the protocol of Floková and co-workers (2014) with modifications. Namely, a StrataX 30 mg/3 ml SPE-column (Phenomenex) was used. For detection and quantification by liquid chromatography-tandem mass spectroscopy (UPLC-TQS/MS), sample residues were dissolved in 100 µl acetonitrile /water (20:80 v/v) and filtered through a 0.45 mm Minisart SRP4 filter (Sartorius, Goettingen, Germany). Analysis of SA, JA, (−)-JA-Ile and (+)-7-iso-JA-Ile was quantified by comparing retention times and mass transitions with those of unlabeled standards ((−)-JA, (−)-JA-Ile, and SA) using a Waters XevoTQS mass spectrometer equipped with an electrospray ionization source coupled to an Acquity ultraperformance liquid chromatography (UPLC) system (Waters, Milford, OH, USA).

Chromatographic separations were conducted using acetonitrile/water (+0.1% formic acid) on an Acquity UPLC BEH C18 column (2.1 mm × 100 mm, 1.7 µm, Waters) at 40°C with a flowrate of 0.25 ml/min. The column was equilibrated for 30 min using acetonitrile/water (20:80 v/v) + 0.1% formic acid. Samples were analyzed by injecting 5 µl, followed by an elution program of 17 min in which the acetonitrile fraction linearly increased from 20% (v/v) to 70% (v/v). The column was washed after every sample by increasing the acetonitrile fraction to 100% in 1 min and maintaining this concentration for 1 min. The acetonitrile fraction was reduced to 20% in 1 min and maintained at this concentration for 1 min before injecting the next sample.

The capillary voltage was set at 2.5 kV, the source temperature at 150 °C and the desolvation temperature at 500°C. Cone and desolvation gas flows were set to 150 and 800 l h^−1^, respectively. The cone voltage was set to 40 eV. Multiple reaction monitoring (MRM) was used for quantification (Table **S2**). Parent-Daughter transitions and cone voltages were optimized for each standard compound using the IntelliStart MS Console (Waters, Milford, OH, USA). Argon was used for fragmentation by collision-induced dissociation. To determine sample concentrations, a 5-point calibration curve was constructed for each compound ranging from 0.1 µM to 19 pM, in addition to a known amount of an appropriate deuterium-labelled internal standard. Peak quantification was processed with Targetlynx (Waters, Milford, OH, USA). Samples were normalized for the internal standard recovery and sample weight. All values were normalized to the 0 dpi control timepoint.

## Results

### Dynamic transcriptional changes in roots upon foliar infection by *Hpa*

To investigate root responses involved in SBL formation, we analyzed transcriptional changes in Arabidopsis Col-0 roots following *Hpa* infection of the leaves. Approximately thirty seedlings per pot were grown in Reijerscamp field soil (Berendsen *et al*. 2018). When two weeks old, the leaves were sprayed with either a *Hpa* spore suspension or water (mock). Root samples from these conditioning-phase plants were collected at seven consecutive time points (1-7 dpi), with each individual pot serving as a biological replicate and for each time point and treatment the RNA from 4 to 6 biological replicates was sequenced (Fig. **1a**). To verify that SBL was successfully established in this experiment, a subset of the pots was subjected to an SBL assay. For this, new Col-0 seeds were sown in soil from which the shoot tissue of the conditioning-phase plants had been removed at 7 dpi. Two weeks later, all these response-phase seedlings were inoculated with *Hpa*, and disease severity was assessed one week post inoculation. At this time point, *Hpa* spore counts were significantly lower in seedlings grown on *Hpa*-conditioned soil compared to mock-conditioned soil (Fig. **1b**), confirming that *Hpa* infection during the conditioning phase successfully induced a disease-suppressive SBL in this experiment.

**Fig. 1.**
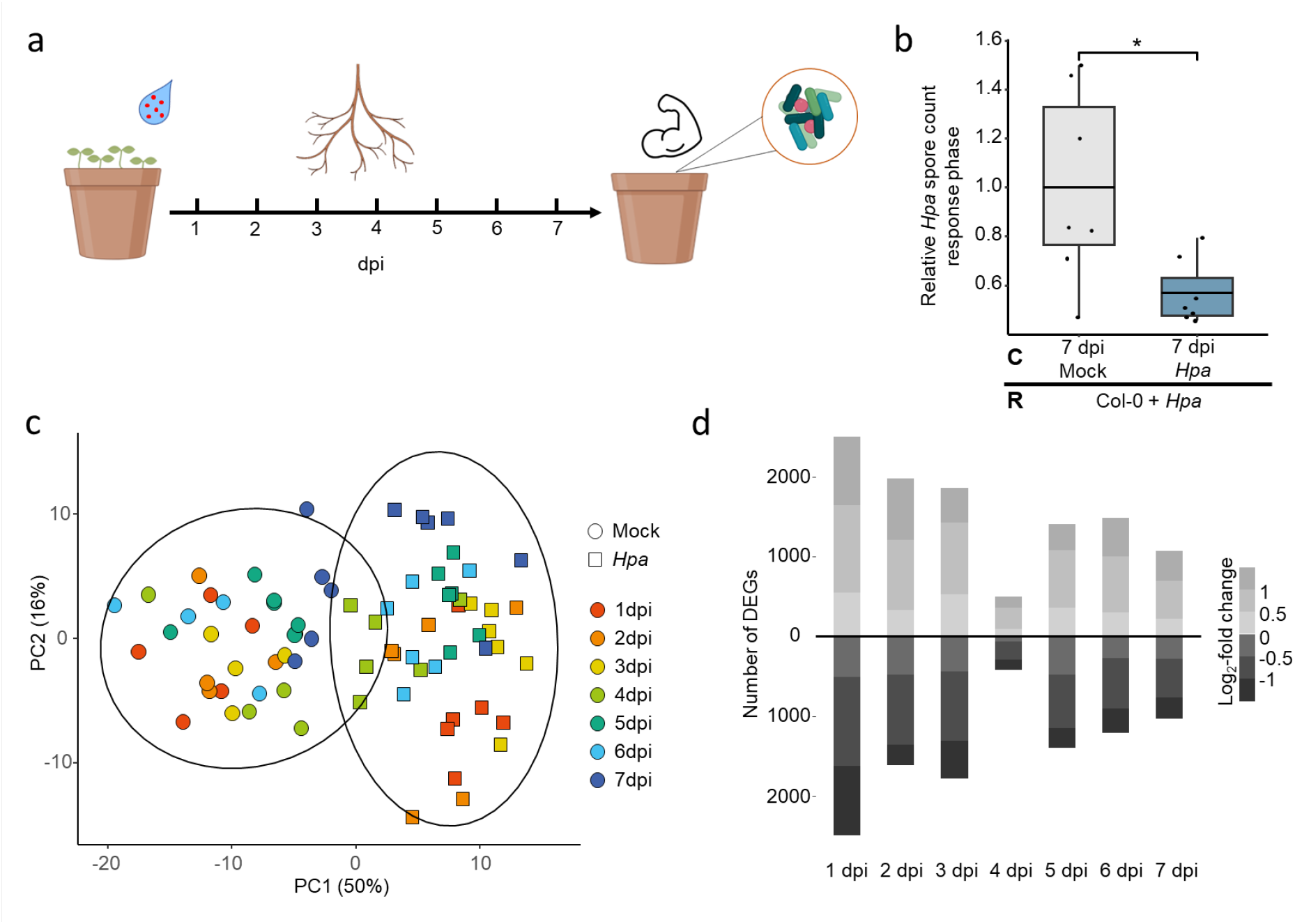
Description of the time-series transcriptome experiment on roots of foliar *Hpa*-infected Arabidopsis. (a) Setup of 7-day time-series RNA-seq experiment. Roots were harvested at 7 consecutive days from pots containing 2- to 3-week-old Arabidopsis seedlings, grown in natural soil, that were leaf-treated with *Hpa* spores or water (mock). (b) Spore production of *Hpa* in response (R) populations of Arabidopsis Col-0 plants growing in soil conditioned (C) by populations of Col-0 plants that were either mock treated or inoculated with *Hpa* for 7 days, following the same procedure as used for the RNA-seq experiment. Relative *Hpa* spore counts were normalized to the mock-treated group in the condition phase for each time point. Asterisks indicate statistically significant differences between mock and *Hpa* treatment within each time point (one-tailed Student’s t-test; * *P* < 0.05, ** *P* < 0.01, *n* = 10-12 pots per experiment). (c) PCA analysis based on the normalized read counts of all genes sequenced from the roots of plants that were leaf treated with *Hpa* or mock at different timepoints. (d) Number of differentially expressed genes (DEGs) (up- or downregulated) in *Hpa* samples compared to mock samples at each timepoint. The bars categorize gene expression changes based on the level of the log_2_-fold changes (*Hpa*/mock), which are depicted as different shades of grey.

The RNA-seq analysis of roots revealed distinct transcriptomic changes in response to *Hpa* infection. Principal component analysis (PCA) indicated that treatment type (mock vs *Hpa*) was the main source of variation, accounting for 50% of the variance along the first principal component (PC1) (Fig. **1c**). Replicate samples clustered closely, indicating low biological variability within each treatment and time point. Pairwise comparisons between *Hpa*- and mock-treated samples at each time point identified DEGs, with approximately equal numbers of genes upregulated and downregulated (Fig. **1d**; for normalized read counts and DEGs profile see Dataset **S1** and **S2**). Strikingly, substantial transcriptomic reprogramming occurred as early as 1 dpi, with nearly 5,000 DEGs detected. Although the number of DEGs gradually declined over time, more than 2,000 DEGs remained at 7 dpi, indicating a sustained transcriptional response in roots to foliar *Hpa* infection.

We next investigated the temporal co-expression patterns of DEGs. A GLM analysis was performed using DESeq2 on the time-series transcriptome data, incorporating time and treatment as factors. This analysis identified 7,498 genes with significantly altered expression in response to *Hpa* infection over time. To explore expression dynamics, we analyzed the coherence of these 7,498 DEGs using SplineCluster, which grouped them into 28 clusters (Fig. **S2**), and Dynamic Regulatory Events Miner (DREM), which identified 20 regulatory paths (Fig. **2a**). The majority of clusters and paths showed continuous up- or downregulation, starting as early as 1 or 2 dpi, while some others displayed transient early or late expression changes. These findings demonstrate that *Hpa* infection generally induces a significant and sustained transcriptional reprogramming in roots, with distinct temporal expression patterns emerging across the course of the infection.

**Fig. 2.**
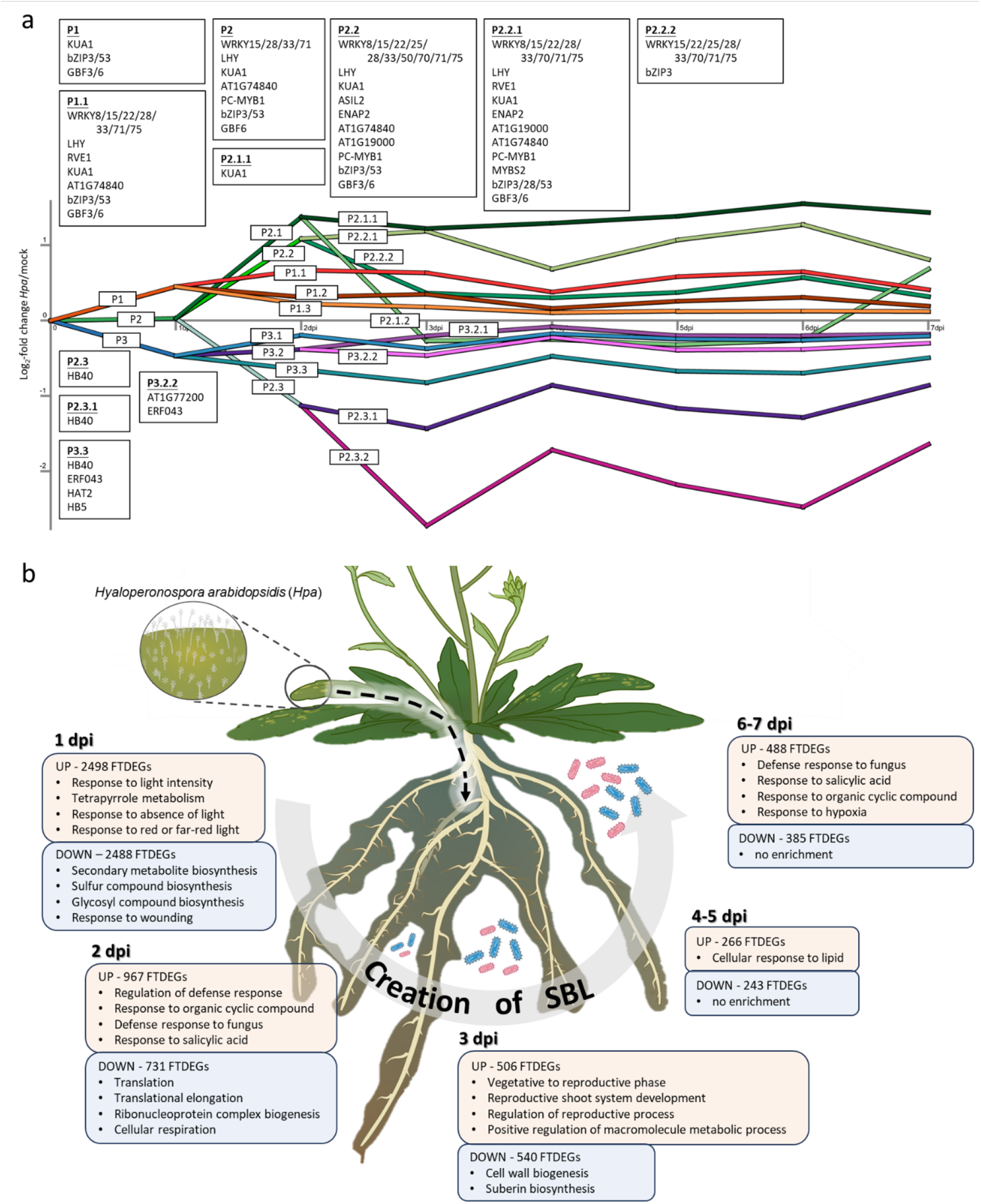
Temporal roles of TFs and biological processes in the root GRN induced by foliar infection *Hpa* infection. (a) Dynamic GRN model. A total of 7498 DEGs over the 7-day time course following *Hpa* inoculation were clustered into 20 distinct paths using Dynamic Regulatory Events Miner (DREM), based on log_2_-fold changes between *Hpa-* and mock-treated plants). TFs significantly associated with the regulation of DEGs within each path were as identified via transcription factor binding site enrichment analysis. In this figure only the most important 36 TFs are shown, which fit all the transcription factor binding site selection criteria (see Fig. 3 for how they were selected). The full list of enriched TFs per path can be found in Dataset **S16** (method SEA) and **S18** (method DAP-seq). (b) GO term enrichment for genes that were differentially expressed for the first time (FTDEG) per time point. A representative selection of all timepoints is depicted. FTDEGs are categorized by upregulation (UP) and downregulation (DOWN). The four most significantly enriched GO terms are shown for each category.

To gain insight into the biological processes in roots influenced by foliar *Hpa* infection, we performed GO term enrichment analysis on DEGs at each time point. The GO term “Response to light intensity” was upregulated as early as 1 dpi and remained elevated until 7 dpi (Fig. **S3**, Dataset **S3-9**). In contrast, biosynthesis of secondary metabolites, including glucosinolates and phenylpropanoids, was rapidly repressed within 1 day and remained downregulated throughout the time course, suggesting their importance in *Hpa*-induced root responses. However, individual gene expression patterns were more dynamic, as shown by SplineCluster and DREM analyses (Fig. **S2** and Fig. **2a**), highlighting the need for a fine-grained temporal analysis. To determine the onset of transcriptional change at the gene level, we identified the first time point of differential expression of each gene (FTDEG) (Dataset **S10**, Fig. **2b**). Over 80% of DEGs were already differentially expressed in roots within 3 days after leaf inoculation, demonstrating that the root transcriptional response to *Hpa* infection is very rapid. Nevertheless, newly regulated genes continued to emerge over time, potentially reinforcing earlier responses or initiating additional biological processes. GO enrichment analysis of FTDEGs provided deeper insights into the temporal sequence of biological responses and the roles of newly activated genes (Fig. **2**). Notably, defense responses were activated at 2 dpi, coinciding with the induction of the SA-responsive WRKY TF genes *WRKY7/15/18/33/51/54* (Dataset **S11-15**). At 7 dpi, a second wave of defense-related gene activation was observed, which included *NPR1* and additional WRKY genes including *WRKY8/31/40/70*. Conversely, GO terms such as "secondary metabolites biosynthesis" and "response to light intensity" were no longer enriched in FTDEGs after 1 dpi, suggesting that most genes in these categories were induced or repressed early and maintained their expression levels throughout the experiment. Overall, the transcription of 30 different WRKY genes was enhanced upon *Hpa* infection, highlighting their extensive involvement in the root’s transcriptional response and their potential role in conditioning the soil for a disease-suppressive legacy.

### Regulators of root responses upon *Hpa* infection

TFs play a crucial role in regulating transcriptional reprogramming. Specific TFs are likely to be essential for SBL formation, acting as regulators of key modules in the GRN activated in roots following *Hpa* infection. To identify candidate TFs, we analyzed DNA sequence motifs enriched in the promoter regions of genes from co-expression clusters and regulatory paths identified by SplineCluster and DREM analyses (Fig. **S2** and Fig. **2a**). We first performed SEA analysis using MEME Suite, scanning 568 Arabidopsis TF motifs from the JASPAR database for motif enrichment based on motif counts. This analysis identified 356 significantly overrepresented TF motifs across the clusters and paths (Dataset **S16**, **S17**). As a complementary strategy, we performed transcription factor binding site enrichment analysis using publicly available DAP-seq data for 347 TFs (O’Malley *et al*. 2016; Aerts *et al*. 2024), which identified 117 overrepresented TFs (Fig. **S4**, Dataset **S18**, **S19**). There was an overlap of 65 TFs between the SEA and DAP-seq analyses (Dataset **S20**). To refine this list, we further filtered for TFs that were themselves differentially expressed in the GLM analysis, resulting in a set of 36 candidate TFs (Dataset **S21**). These 36 TFs were considered the most reliably predicted key regulators (Fig. **3a**). Notably, 10 of these TFs belong to the WRKY family, consistent with the FTDEG analysis, which identified multiple WRKY genes upregulated starting at 2 dpi and continuing through 7 dpi. The prominence of WRKYs was also supported by the DREM analysis, which identified them as predicted regulators of upregulated paths throughout the time course (Fig. **2a**), further highlighting their potential role in mediating root responses and contributing to SBL formation. Next, we assessed the relationship among the 36 candidate TFs assessing the similarity in their predicted binding to the 7,498 DEGs identified in the roots of *Hpa*-infected plants. Based on DAP-seq data, 5,882 of these DEGs were predicted to be targets of at least one of the 36 TFs (Fig. **3b**). Notably, several TFs families, particularly WRKYs and bZIPs, displayed substantial overlap in their predicted target genes (Fig. **3c**). Interestingly, most WRKY TFs exhibited increased expression, along with their predicted target genes, suggesting coordinated activation. In contrast, the bZIP TFs *GBF3* and *GBF6* showed divergent expression patterns: *GBF3* was upregulated, while *GBF6* was downregulated. Despite this, they shared a large set of predicted target genes, the majority of which were upregulated. A similar discrepancy was observed for other TFs: *HB40* was upregulated, while its predicted targets were predominantly downregulated, whereas *LHY* showed the opposite pattern, being downregulated while its predicted targets were upregulated (see also Fig. **2a**).

**Fig. 3.**
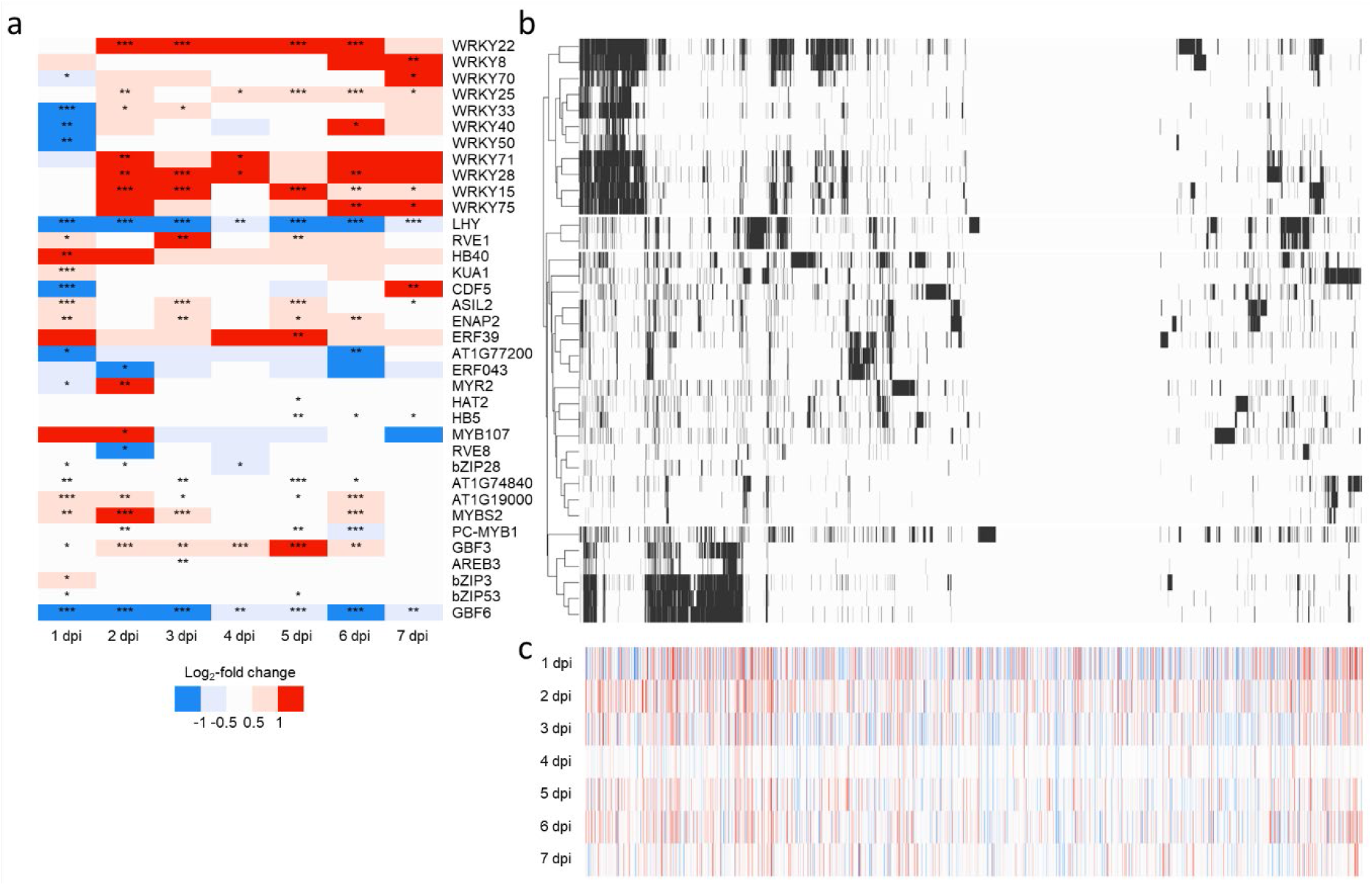
Predicted regulators of the root transcriptional responses to foliar *Hpa* infection. The three plots are arranged to share axes for better visual comparison: specifically, (a) and (b) share the y-axis, which lists TFs; (b) and (c) share the x-axis, representing DEGs identified by GLM analysis. Due to spacing constraints, DEG names are not shown on the figure. (a) Expression profiles of TF genes whose predicted target genes are significantly differentially expressed. The heatmap shows the log_2_-fold change in expression in roots between *Hpa*-infected and mock-treated plants across 7 time points following *Hpa* infection. Each row represents a TF gene associated with enriched transcription factor binding sites in its target genes, as depicted in (b). Asterisks indicate significantly differential expression based on DESeq2 analysis (* *P* < 0.05, ** *P* < 0.01, *** *P* < 0.001). (b) Heatmap showing the interaction between the TFs depicted in (a) with the 7498 DEGs identified in *Hpa*-induced roots using GLM analysis. Transcription factor binding site enrichment was determined using filtered DAP-seq data. (c) Expression profiles of the 7498 DEGs identified in *Hpa*-induced roots by GLM analysis. Values represent log_2_-fold change between *Hpa* and mock treatments across 7 time points following *Hpa* infection. Each column represents one DEG, for which enrichment of transcription factor binding sites is shown in (b).

### Tryptophan-derived defense compounds and flavonoids are important SBL formation factors

The GO term “glucosinolate biosynthetic process” was among the most significantly enriched categories, with over 60% (28/45) of annotated genes in this pathway exhibiting significant transcriptional changes during the time course. Overall, genes encoding enzymes and TFs involved in both the AGS and IGS biosynthesis pathways were downregulated in roots following *Hpa* infection (for IGS genes see Fig. **4a**; overview of both IGS and AGS genes in Fig. **S5**). However, a notable exception was observed for the four genes encoding indole glucosinolate O-methyltransferase (*IGMT1-4*), which catalyze the conversion of 4OH-I3M to 4MO-I3M (Pfalz *et al*. 2011). These genes exhibited a robust increase in transcript levels, up to 7-fold higher in *Hpa*-infected samples compared to mock-treated controls (Fig. **4a,b**). In contrast, genes involved in the synthesis of 1MO-I3M, which competes with 4MO-I3M for the same precursor I3M, were downregulated. These findings suggest that *Hpa* infection selectively reprograms glucosinolate biosynthesis in roots, with transcriptional activation of the 4MO-I3M branch despite the overall repression of glucosinolate biosynthesis.

**Fig. 4.**
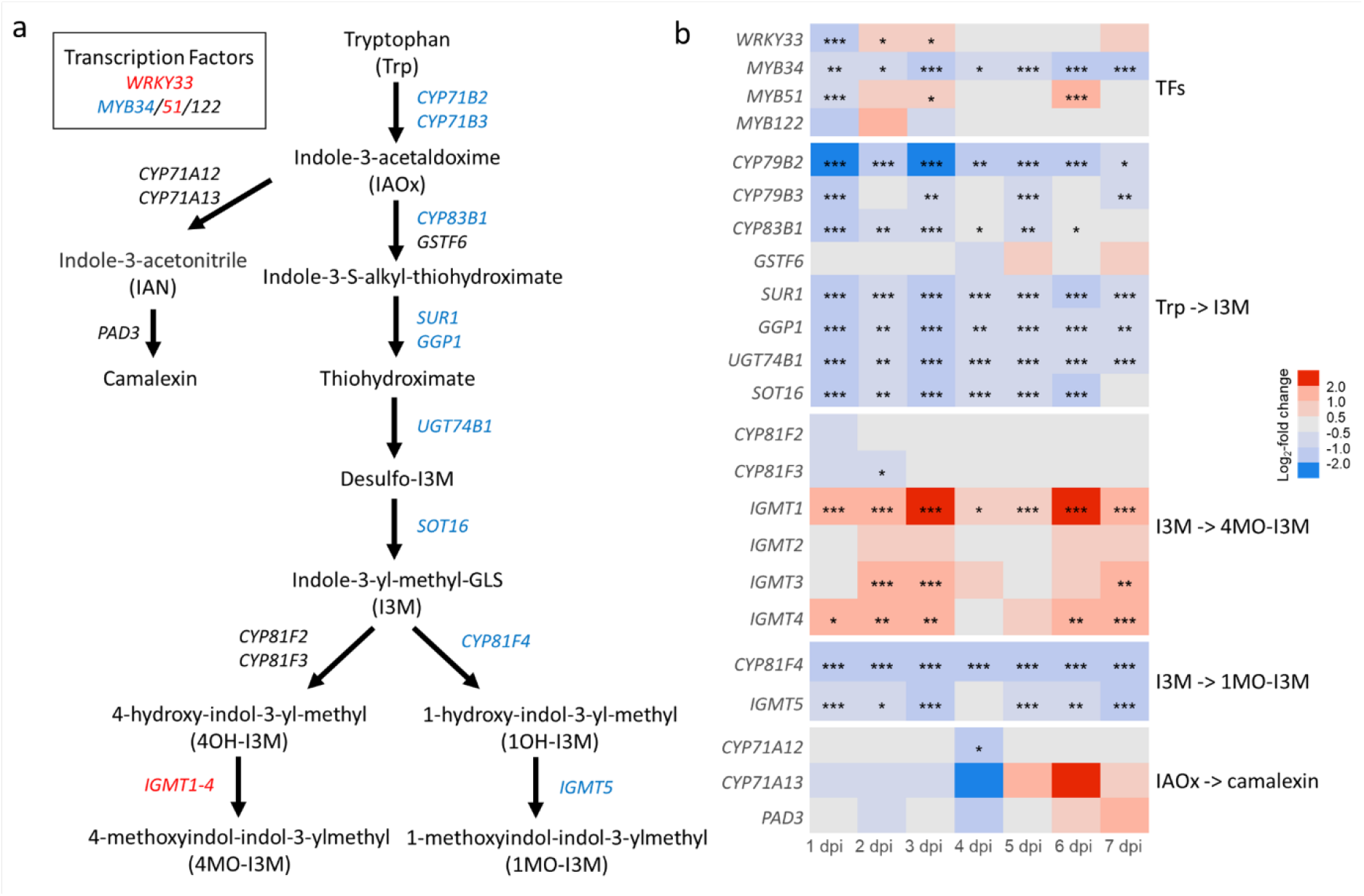
Changes in transcript levels of genes from the biosynthesis pathways of tryptophan-derived defense compounds in roots following foliar *Hpa* infection. (a) Detailed biosynthesis pathways of tryptophan-derived IGS (Pfalz *et al*. 2011) and camalexin. Compounds are indicated in black straight letters.Genes encoding transcription factors are shown within the box, whereas genes encoding enzymes are indicated in italics along the arrows. Gene names are colored in red, blue, or black depending on their average change in expression level over time, namely upregulated, downregulated, or not changed, respectively. (b) Temporal expression profiles of genes in the IGS and camalexin biosynthesis pathways following *Hpa* inoculation. Rows indicate the log_2_-fold change between *Hpa* and mock treatments at 1-7 dpi, following the color range indicated in the legend. Asterisks indicate significantly different expression based on DESeq2 analysis (* *P* < 0.05, ** *P* < 0.01, *** *P* < 0.001).

To investigate the biological significance of the general repression and selective upregulation of glucosinolate biosynthesis genes, we examined the ability of Arabidopsis mutants impaired in key biosynthetic (sub-)pathways to establish a SBL. Mutant *myb28/29* accumulates low levels of AGS (Sønderby *et al*. 2010); *cyp81f4* is deficient in the production of 1MO-I3M (Pfalz *et al*. 2011); and *cyp81f2* exhibits reduced 4MO-I3M production due to impaired conversion from I3M to its precursor 4OH-I3M (Pfalz *et al*. 2009). Given that WRKY33 was consistently identified across our analyses as a key regulatory TF, and is known to regulate 4MO-I3M biosynthesis (Tao *et al*. 2022) and also controls camalexin production, another S-containing antimicrobial compound that is also derived from tryptophan (Birkenbihl *et al*. 2012), we included the *wrky33* mutant and the camalexin biosynthesis mutant *pad3* in our SBL assay. Our results revealed that WRKY33 and the biosynthesis pathways of 4MO-I3M and camalexin are crucial for *Hpa*-induced SBL formation. This was evidenced by the absence of enhanced protection in wild-type Col-0 plants grown on soil previously conditioned by *Hpa* infection of these mutants (Fig. **5a**). Notably, wild-type plants grown on soil conditioned by mock treatment of the *wrky33* and *cyp81f2* mutants already exhibited reduced sensitivity to *Hpa* infection (Dataset **S21**), suggesting that their constitutive alterations in immune-homeostasis led to a protective soil, via distinct mechanisms than *Hpa*-induced SBL (Fig. **5a**). Two-way ANOVA statistical analysis, which we performed on all mutant bioassays in this study (Dataset **S21**), revealed that *Hpa* infection significantly reverted the enhanced protection of the mutants, corroborating interplay between the different mechanisms. In contrast to WRKY33 and the biosynthesis pathways of 4MO-I3M and camalexin, our results indicate that neither IGS 1MO-I3M, produced via CYP81F4, nor AGS, regulated by MYB28/29 is required for SBL establishment. This is demonstrated by the enhanced protection that was still evident in the response phase of wild-type plants grown in soil conditioned by *cyp81f4* or *my29/29* mutants (Fig. **5a**).

**Fig. 5.**
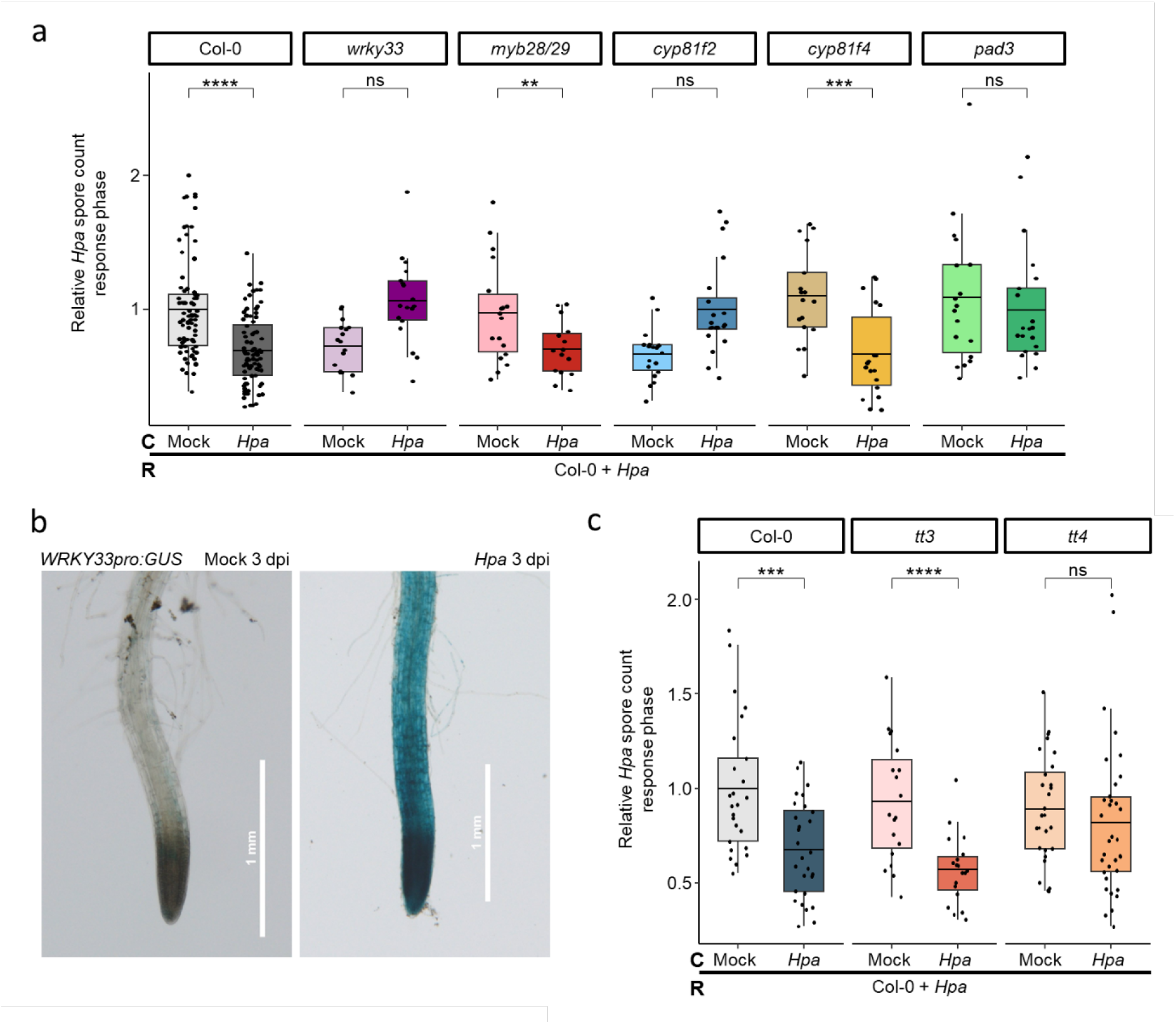
Biosynthesis genes of tryptophan-derived defense compounds and flavonoids are important SBL formation factors. (a) Spore production of *Hpa* in response (R) populations of Col-0 plants growing in soil conditioned (C) by populations of wild-type Col-0 or mutant lines (*wrky33*, *myb28/29*, *cyp81f2*, *cyp81f4*, and *pad3*) that were either mock treated or inoculated with *Hpa* for 7 days. Relative *Hpa* spore count represents the combined results of nine experiments, with spore counts in each experiment normalized to the mock-treated group in the condition phase (one-tailed Student’s t-test; ** *P* < 0.01, *** *P* < 0.001, **** *P* < 0.0001; ns, non-significant; *n* = 10-12 pots per experiment). (b) GUS staining of reporter line *WRKY33pro:GUS* under mock- and *Hpa*-treated conditions at 3 dpi. The pictures show representative tips of lateral roots grown *in vivo* in the “tea bag system” (see Fig. **S1**). For quantification, see Fig. **S6**. (c) Spore production of *Hpa* in response (R) populations of Col-0 plants growing in soil conditioned (C) by populations of wild-type Col-0 or mutant lines (*tt3* and *tt4*) that were either mock treated or inoculated with *Hpa* for 7 days. Results represent the combined data from two independent experiments. See legend of (a) for more details.

WRKY33, a multifaceted defense response regulator, plays a crucial role in the establishment of *Hpa*-induced SBL, as indicated by its enrichment in our transcription factor binding site analysis and the inability of the *wrky33* mutant to generate SBL. To better understand its spatial expression in root tissue following foliar *Hpa* infection, we visualized its expression pattern using a *WRKY33pro:GUS* reporter line. To isolate clean and intact Arabidopsis root tissue grown in natural soil, we developed the "tea bag system" of which the setup is described and visualized in detail in Materials and Methods (Fig. **S1a,b**). This system consists of a nylon mesh pocket in which the root system grows, and which is placed inside a pot filled with Reijerscamp field soil. While the mesh prevents penetration of the roots into the soil, it allows the exchange of microbes and metabolites between compartments. Given this physical separation, we validated that the system retains the capacity of plants to generate an SBL (Fig. **S1c**). Interestingly, upon *Hpa* infection, the *WRKY33pro:GUS* line showed GUS staining in most root tips, including both the primary and lateral roots, indicating strong induction of *WRKY33* (Fig. **5b**). In contrast, fewer root tips were stained in the mock-treated group (quantified in Fig. **S6**), and the staining was much lighter (Fig. **5b**). This result aligns with our transcriptome data, which showed overall *Hpa*-induced upregulation of *WRKY33* after an initial downregulation at 1 dpi (Fig. **4b**). The GUS staining data further show that the transcriptional regulation is primarily localized to root tips, where WRKY33 may influence multiple downstream responses, including tryptophan-derived defenses.

Our transcriptome analysis revealed a downregulation of the phenylpropanoid biosynthesis pathway, particularly genes involved in flavonoid biosynthesis, such as *TT4* and *UGT78D1,* showed reduced expression. To examine the role of flavonoids and anthocyanins in SBL formation, we tested two biosynthesis mutants, *tt4* and *tt3*. *TT4* encodes chalcone synthase, the first enzyme in the flavonoid biosynthesis pathway, while *TT3* encodes dihydroflavonol 4-reductase, which catalyzes the conversion of dihydroquercetin to leucocyanidin, the initial step in the flavonoid-to-anthocyanin branch. Although *TT3* transcript levels were undetectable in roots, we speculated that its metabolic products might originate in the shoots and be transported to the roots for secretion into the soil. This hypothesis is supported by previous findings that certain (undefined) phenolic compounds were reduced in the root exudates of *Hpa*-infected plants (Vismans *et al*. 2022). Our bioassay results showed that *tt4* failed to establish SBL, whereas *tt3* retained this ability (Fig. **5c**). These findings suggest that flavonoid biosynthesis is essential for SBL formation, while anthocyanins and other downstream secondary metabolites synthesized via TT3 are not required.

### Involvement of JA and SA pathways in SBL formation

The general downregulation of genes involved in glucosinolate and phenylpropanoid biosynthesis (e.g., coumarins and flavonoids) suggests that the JA pathway, known to promote the production of these compounds, may be repressed. However, genes typically responsive to JA signaling, such as those encoding LOXs, JAZs, MYCs and ERFs, were not significantly downregulated, indicating that *Hpa*-induced effects on the JA pathway in roots may be subtle. To investigate the role of the JA pathway in SBL formation, we tested the effect of TF MYC2, a central regulator of JA responses that links JA signaling to the transcriptional control of glucosinolate and flavonoid biosynthesis (Schweizer *et al*. 2013; Wang *et al*. 2021a), two metabolic pathways prominently affected in our transcriptome study. As shown in Fig. **6a**, unlike Col-0 wild-type plants, *myc2* mutants infected by *Hpa* in the conditioning phase, did not significantly increase *Hpa* resistance in the response population of Col-0 plants. This suggests that MYC2 is required for SBL formation mediated by *Hpa* infection, possibly via dynamic inhibition of MYC2 activity.

**Fig. 6.**
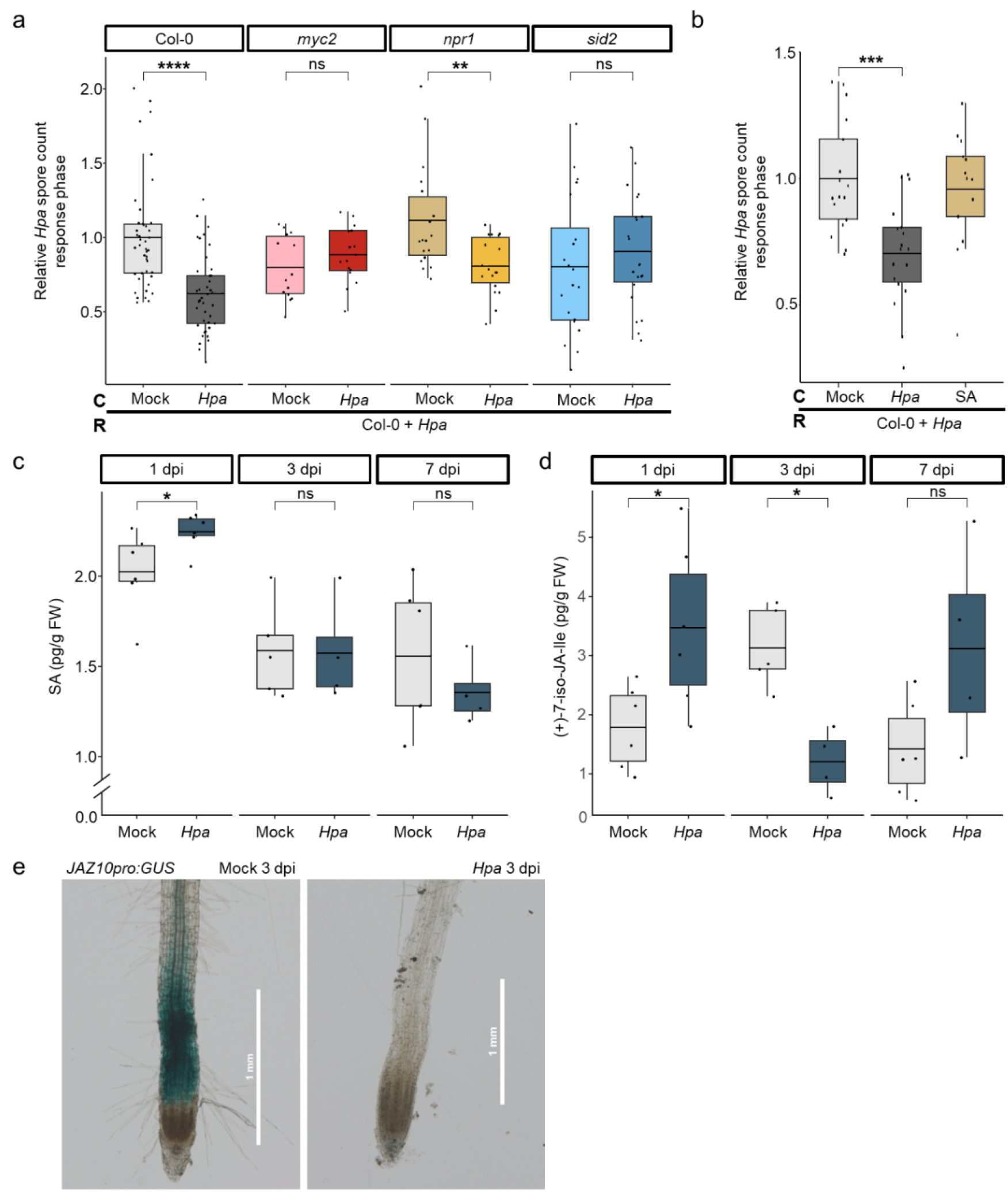
Involvement of SA and JA pathways in SBL formation by *Hpa* foliar infection. (a) Spore production of *Hpa* in response (R) populations of Col-0 plants growing in soil conditioned (C) by populations of wild-type Col-0 or mutant lines (*myc2, npr1* and *sid2*) that were either mock treated or inoculated with *Hpa* for 7 days. Spore production of *Hpa* on Col-0 plants in the response phase at one week after inoculation. Relative *Hpa* spore count represents the combined results of five experiments, with spore counts in each experiment normalized to the mock-treated group in the condition phase. Asterisks indicate statistically significant differences (one-tailed Student’s t-test; ** *P* < 0.01,*** *P* < 0.001, **** *P* < 0.0001; ns, non-significant; *n* = 10-12 pots per experiment). (b) Spore production of *Hpa* in response (R) populations of Col-0 plants growing in soil conditioned (C) by populations of Col-0 plants that were either treated with *Hpa*, mock or 1 mM SA for 7 days. Results represent the combined data from two independent experiments. See legend of (a) for more details. (c) and (d) Concentrations of the hormones SA and (+)-7-iso-JA-Ile in roots at 1, 3 and 7 dpi after foliar mock or *Hpa* treatment. Hormone levels are normalized to fresh weight (FW) of each sample. Asterisks indicate statistically significant differences (Student’s t-test; * *P*<0.05; *n* = 6). (e) GUS staining of reporter line *JAZ10pro:GUS* in primary root tips under mock- and *Hpa*-treated conditions at 3 dpi. The pictures show representative tips of primary roots grown *in vivo* in the “tea bag system” (see Fig. **S1**).

Meanwhile, our GO enrichment analysis indicated that the pathway of the other major defense hormone, SA, could be activated in roots. SA production has previously been demonstrated to be crucial for resistance to *Hpa* (Delaney *et al*. 1994). Therefore, we assessed its role in *Hpa*-induced SBL by performing bioassays with the *sid2* mutant, which is deficient in SA biosynthesis, and the *npr1* mutant, which is non-responsive to SA. Interestingly, *sid2* failed to generate SBL, whereas *npr1* was still able to effectively condition the soil upon *Hpa* infection, leading to enhanced *Hpa* resistance in the subsequent generation of Col-0 plants (Fig. **6a**). We also applied SA exogenously to the leaves of wild-type plants; however, this treatment did not trigger SBL, as the response population was as susceptible to *Hpa* as those conditioned with mock-treated plants (Fig. **6b**). This suggests that SA biosynthesis is required for SBL formation upon *Hpa* infection, but that SA treatment alone is not sufficient to mimic this response, and that NPR1 is not required for *Hpa*-induced SBL.

To investigate changes in the levels of the hormones SA and JA, root and shoot tissues were analyzed at 1, 3 and 7 days after *Hpa* inoculation or mock treatment of the leaves. In roots, the SA levels were significantly elevated at 1 dpi but remained unchanged at later time points (Fig. **6c**), whereas in leaves, SA levels increased progressively throughout the course of infection (Fig. **S7a**). JA and its bioactive derivative, (+)-7-iso-JA-Ile (Staswick and Tiryaki 2004), were elevated in roots at 1 dpi but showed a significant decrease at 3 dpi (Fig. **S7b**, Fig. **6d**). In contrast, both JA and (+)-7-iso-JA-Ile levels in the leaves increased only at 7 dpi, coinciding with visible leaf surface damage associated with *Hpa* sporulation (Fig. **S7c,d**). These early and transient changes in SA and JA levels at 1 dpi corresponded with the transcriptomic data, which showed the highest number of DEGs at this time point (Fig. **1d**). The increase in SA levels was followed by upregulation of SA-related genes at 2 dpi, as described above (Fig. **2b**, Dataset **S4** and **S12**). While the reduction in JA levels at 3 dpi coincided with downregulation of fatty acid biosynthesis genes (Dataset **S5** and **S13**), the transient increase in JA at 1 dpi did not appear to trigger extensive transcriptional responses. Notably, the suppression of secondary metabolite biosynthetic pathways, observed at 1 dpi (Dataset **S3** and **S11**), preceded the decline in JA accumulation at 3 dpi, suggesting that this repression was unlikely to be directly mediated by JA.

To further elucidate the role of JA, we employed the afore-mentioned “tea bag system” to test JA responsiveness in soil-grown roots following *Hpa* infection, using GUS reporter line *JAZ10pro:GUS*. JAZ10, a negative regulator of JA signaling, which serves as a transcriptional marker for JA pathway activity in plant tissues (Chung *et al*. 2008). In mock-treated plants, we detected constitutive GUS staining in the root tips of *JAZ10pro:GUS* seedlings. However, at 3 dpi with *Hpa*, this GUS activity was nearly abolished (Fig. **6e**). Even after exogenous MeJA application to the roots, *JAZ10pro:GUS* plants infected with *Hpa* displayed a reduced GUS signal in the root tips in comparison to the enhanced GUS activity in mock-treated plants (Fig. **S8**). These observations are consistent with the significantly reduced JA levels detected at 3 dpi (Fig. **4c**) and support the conclusion that *Hpa* infection represses JA signaling in the roots. Together, these findings confirm that both SA and JA signaling pathways are crucial for in *Hpa*-induced SBL, while also highlighting their complex and context-dependent regulation of induced root responses. Furthermore, our GUS activity assays using *JAZ10pro* and *WRKY33pro* reporters demonstrate that the transcriptional regulation is particularly active in root tips, reinforcing the notion that this region may play an important role in coordinating root responses associated with SBL establishment following *Hpa* infection.

## Discussion

The formation of disease-suppressive soils through microbiome recruitment has been recognized for decades (Schroth and Hancock 1982; Gómez Expósito *et al*. 2017; Spooren *et al*. 2024; Spooren *et al*. 2026), and considerable attention has been given to the roles of individual beneficial microbes, such as *Pseudomonas* spp. and arbuscular mycorrhizal fungi, in elicitation of ISR (Cameron *et al*. 2013; Pieterse *et al*. 2021). However, the plant-driven molecular and metabolic mechanisms underlying selective recruitment of protective microbial consortia leading to disease-suppressive soils remain poorly understood (Yuan *et al*. 2018; Wang *et al*. 2021b). In this study, we uncovered a root-specific transcriptional program activated in Arabidopsis following foliar infection by *Hpa* that underpins the formation of a disease-suppressive soil that enhances resistance to *Hpa* in successive plant populations grown in the same soil. Through time-series RNA-seq and GRN analysis, we demonstrate that root transcriptional reprogramming begins as early as 1 dpi and persists over a seven-day infection window, ultimately facilitating SBL and enhancing resistance in subsequent plant generations. Our data supports a model (Fig. **7**) in which *Hpa* infection in the shoot triggers SA biosynthesis and canonical defense responses. In parallel, yet-unknown mobile signals induce root-localized responses, including early SA accumulation and repression of JA signaling and secondary metabolite biosynthesis, with specific components of these root responses contributing to the subsequent establishment of SBL.

**Fig. 7.**
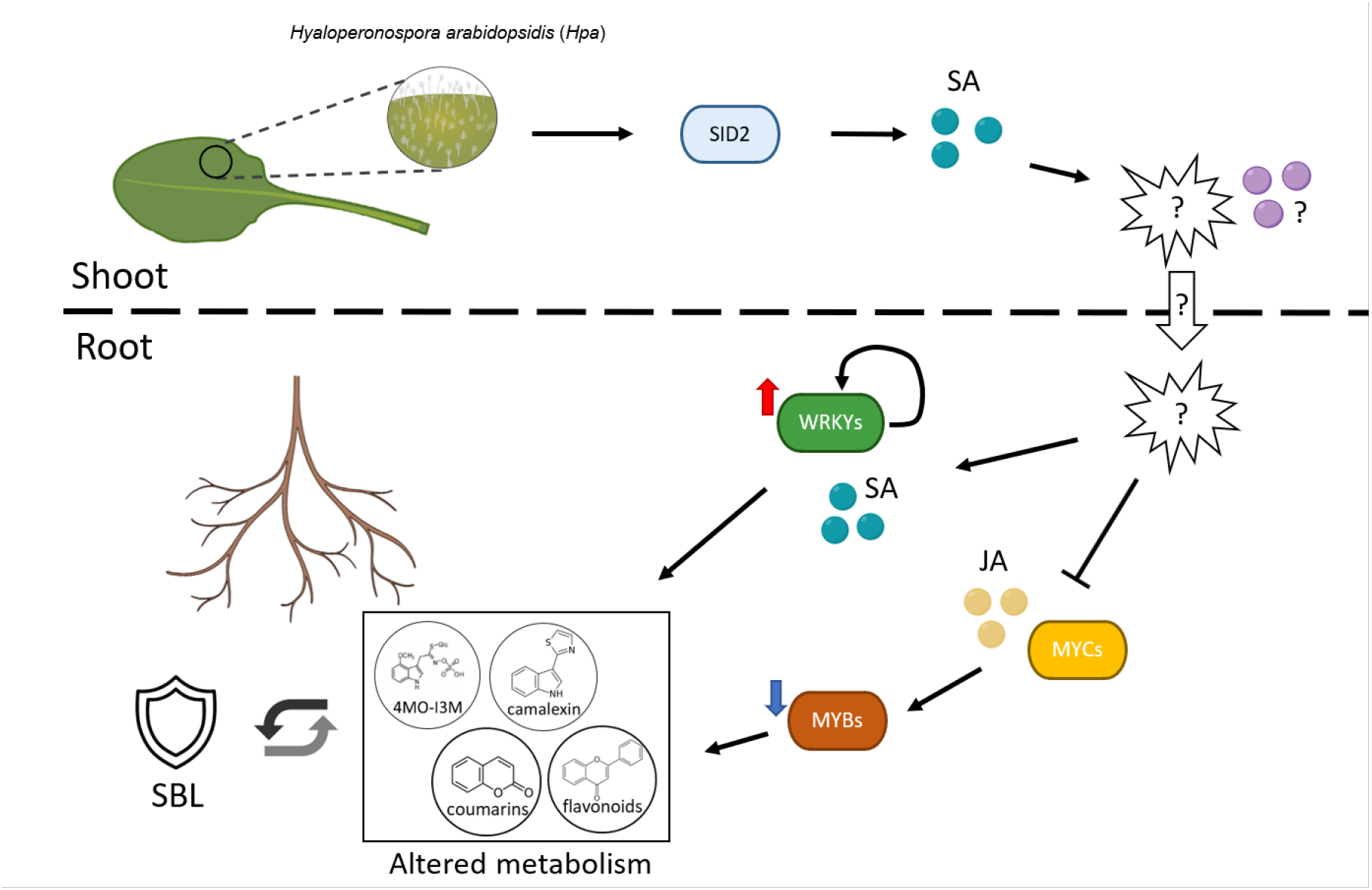
Hypothetical model of how Arabidopsis establishes SBL upon *Hpa* infection. The diagram illustrates key biological processes and regulatory pathways involved in SBL formation. Leaf infection triggers SID2-dependent SA biosynthesis and local defense. Unidentified systemic signals are proposed to communicate the infection status from shoots to roots. In the roots SA signaling is activated and JA signaling suppressed, which is associated with induction of WRKYs and repression of MYBs, accompanied by altered secondary metabolite patterns, including changes in indole- and phenylpropanoid-related pathways. These metabolic changes may subsequently influence the rhizosphere chemical environment and microbial interactions causal to SBL establishment, but their relative contributions remain to be resolved. Black arrows, activation; bar-headed lines, repression; red and blue arrows, gene upregulation and downregulation, respectively; question-mark explosions, unknown signals or steps.

While the role of SA in systemic acquired resistance is well established, our results provide evidence for its role in SBL formation. We show that SA biosynthesis via SID2 is essential in the conditioning phase, whereas SA signaling via NPR1 is dispensable. These findings suggest that SBL formation relies on NPR1-independent responses, possibly involving alternative downstream signaling routes or TFs. This is consistent with prior findings that *sid2* mutant plants more strongly affect rhizosphere microbiome composition than *npr1* mutant plants (Lebeis *et al*. 2015). Hormone profiling revealed an early SA surge in roots at 1 dpi, prior to SA accumulation in shoots (Fig. **6c**, Fig. **S7a**), coinciding with the first wave of SA-responsive gene expression (Fig. **2b**). These rapid root responses likely reflect systemic signaling from the shoot, which is triggered by early pathogen recognition. A second wave of SA-responsive gene expression at 7 dpi, including *NPR1*, *WRKY38*, *WRKY40*, and *WRKY70* (Fig. **S3**), may result from direct root-microbiome interactions or systemic signaling linked to pathogen sporulation, as we found that SBL is established prior to or concurrently with this stage (Fig. **1b**). Notably, exogenous SA application to the leaves failed to induce SBL (Fig. **6b**), underscoring that SA accumulation in the shoot alone is insufficient and suggesting the involvement of *Hpa* infection-related cues or complex hormonal crosstalk. This raises the possibility that the communication between *Hpa*-infected shoots and the roots involves multiple systemic signals rather than a single mobile factor. Long-distance plant responses can involve rapid Ca²⁺/ROS-associated signaling, as well as mobile molecular signals transported between tissues, such as defense-related metabolites, peptides and RNAs (Kehr and Kragler 2018; Kachroo and Kachroo 2020; Gilroy *et al*. 2016). Such signaling mechanisms may contribute to the early and sustained root responses observed here, although identifying the signals involved and their relative contributions to *Hpa*-induced root responses remains an important question for future studies.

JA signaling played a more nuanced role. Although canonical JA-responsive genes were not uniformly downregulated, the JA biosynthesis pathway and several JA-dependent metabolic branches, such as flavonoid and glucosinolate biosynthesis, were suppressed in roots after *Hpa* infection. Surprisingly, the *myc2* mutant, lacking a central transcriptional regulator of JA responses, failed to establish SBL. Thus, despite an overall attenuation of JA signaling, MYC2-dependent transcriptional regulation remains functionally essential for successful *Hpa*-induced soil conditioning. We found a similar pattern in our previous study, where coumarin levels decreased during the conditioning phase, yet the coumarin biosynthesis mutants *f6’h1* and *myb72* were unable to establish SBL (Vismans *et al*. 2022). Together, these findings suggest that MYC2, F6’H1 and MYB72 function as a regulatory switch required to establish the transcriptional program underlying SBL, rather than simply promoting their downstream metabolic pathways. Possibly, the ability to dynamically (down-)regulate these pathways is more important than maintaining high constitutive metabolite levels. Hormone measurements, RNA-seq analyses and GUS reporter assays support a model of transient JA accumulation at 1 dpi, which is followed by a decline (Fig. **6d**), accompanied by an early and overall global repression of secondary metabolism-related gene expression (Fig. **2b**,**4**) and reduced JA signaling in root tips (Fig. **6d,e**). These observations suggest that spatial and dynamic regulation of JA signaling in the roots during foliar *Hpa* infection overrules potential activation by a transient increase in JA level at the initial phase of *Hpa* infection, while the subsequent decline in JA levels may reinforce or stabilize this transcriptional state. How this dynamic regulation of JA signaling and MYC2-controlled metabolic pathways contributes to soil suppressiveness remains an important question for future research.

A key discovery was the selective upregulation of the 4MO-I3M biosynthesis pathway, a branch of indole glucosinolate metabolism regulated by WRKY33 and MYB51, while the 1MO-I3M pathway and AGS biosynthesis were suppressed. Only mutants deficient in 4MO-I3M and camalexin biosynthesis failed to establish SBL, indicating that these tryptophan-derived metabolic pathways play important roles during the SBL conditioning phase. Their involvement may be related to the known antimicrobial and microbiome-associated functions of these metabolites. Consistent with this interpretation, Chroston *et al*. (2024) have shown that disruption of glucosinolate biosynthesis or breakdown pathways significantly alters rhizosphere microbial community composition, enriching or depleting specific bacterial genera, such as *Rhodococcus*, *Flavobacterium*, *Chryseobacterium* and *Caulobacter*. Our findings suggest that 4MO-I3M and camalexin biosynthesis are co-regulated and metabolically linked, and that disruption of one pathway impairs the other. Such interconnectedness underscores the need to study secondary metabolism as an integrated network rather than isolated branches. The complexity in metabolic networks is also reflected in the soil-conditioning phenotypes of *wrky33* and *cyp81f2* mutants. While mock-conditioned soil unexpectedly conferred enhanced resistance (Fig. **5a**, Dataset **S21**), these mutants failed to establish SBL following *Hpa* infection, indicating that their constitutive soil-conditioning phenotypes differ from the SBL response induced by *Hpa* infection. The basis of these contrasting responses remains unclear.

The spatial specificity of *Hpa-*induced gene expression in roots, and particularly in the root tips, further emphasizes their potential importance as hubs for metabolite exudation and microbial interactions during the establishment of SBL. Both *WRKY33pro:GUS* and *JAZ10pro:GUS* reporter lines showed strong activity in root tips upon *Hpa* infection, aligning with prior observations that this region is critical for secretion of defense-related compounds and recruitment of microbiota (Yang and Crowley 2000; DeAngelis *et al*. 2009; Frerigmann and Gigolashvili 2014; Tao *et al*. 2022; Rüger *et al*. 2023).

Our results also refine the role of phenylpropanoid metabolism in SBL. Although this pathway was broadly repressed at the transcriptomic level, we found that flavonoid biosynthesis, rather than anthocyanin production, was required for the establishment of SBL, as demonstrated by the inability of the *tt4* mutant (deficient in flavonoids) but not the *tt3* mutant (deficient in anthocyanins) to establish a protective soil. Given that flavonoids such as apigenin and luteolin have been reported to influence the colonization and enrichment of beneficial taxa including *Oxalobacteraceae*, *Pseudomonas*, and *Gluconacetobacter* (Yu *et al*. 2020; Yu *et al*. 2021; Yan *et al*. 2022), the role of flavonoid biosynthesis in SBL may involve changes in interactions with specific rhizosphere bacteria. These findings suggest that specific phenylpropanoids, likely flavonoids (Fig. **5c**) and coumarins (Vismans *et al*. 2022), contribute to SBL, potentially through their effects on the rhizosphere environment. Together with the roles identified for 4MO-I3M and camalexin, these findings raise the question of how changes in secondary metabolism are translated into changes in the rhizosphere environment during SBL establishment. To get more insights, future studies could combine rhizosphere metabolome analysis with microbiome profiling and functional assays. Exogenous application of selected metabolites, individually or in combination, could provide a complementary approach to assess whether they can influence performance of subsequent plantings either via the microbiome or not.

In conclusion, our study reveals a coordinated root transcriptional program triggered by foliar *Hpa* infection that is essential for the establishment of a disease-suppressive soil. This transcriptional program involves SA and JA signaling, spatially regulated responses in the root tip, and selective activation or repression of secondary metabolism pathways (summarized in Fig. **7**). These findings enhance our understanding of plant-soil interactions and lay the groundwork for future studies investigating how molecular and metabolic processes influence soil states associated with SBL. Integrating transcriptomics with spatially resolved metabolite profiling and microbiome functional analysis will be key to unraveling the intricate plant molecular mechanisms governing SBL formation. In the longer term, understanding these relationships may provide opportunities to manipulate disease-suppressive soil states through selected plant-derived metabolites or microbial communities.

## Acknowledgements

This study was sponsored by the Dutch Research Council (NWO) through the XL program ‘Unwiring beneficial functions and regulatory networks in the plant endosphere’ (grant number OCENW.GROOT.2019.063). We acknowledge the contributions of students Remy Pieterse, Koen Trouw, Laurens Geene, Mieke Brinkman, Judie van Diepen, Stan van Gisbergen, Jana Marx and Duco Heere to experiments in this paper.

## Author contributions

R.Q., R.L.B., C.M.J.P. and S.C.M.V.W. designed the experiments. R.Q., I.G., J.S., X.L. and M.V.L. performed the experiments. F.V. and I.F.K. performed the hormone measurements. R.Q. analyzed the data. R.Q., C.M.J.P. and S.C.M.V.W. wrote the manuscript.

## Data availability

The raw RNA sequencing data generated in this study are available at https://www.ncbi.nlm.nih.gov/sra/PRJNA1310175.

## Supporting Information

**Dataset S1** Normalized read counts of all genes across biological replicates over a 7-day time course following *Hpa* or mock treatment in Col-0 roots.

**Dataset S2** DEG analysis results from time-series RNA-seq of Col-0 roots (*Hpa* vs. mock).

**Dataset S3** GO enrichment of DEGs at 1 dpi (*Hpa* vs. mock) in Col-0 roots.

**Dataset S4** GO enrichment of DEGs at 2 dpi (*Hpa* vs. mock) in Col-0 roots.

**Dataset S5** GO enrichment of DEGs at 3 dpi (*Hpa* vs. mock) in Col-0 roots.

**Dataset S6** GO enrichment of DEGs at 4 dpi (*Hpa* vs. mock) in Col-0 roots.

**Dataset S7** GO enrichment of DEGs at 5 dpi (*Hpa* vs. mock) in Col-0 roots.

**Dataset S8** GO enrichment of DEGs at 6 dpi (*Hpa* vs. mock) in Col-0 roots.

**Dataset S9** GO enrichment of DEGs at 7 dpi (*Hp*a vs. mock) in Col-0 roots.

**Dataset S10** DEG and FTDEG counts.

**Dataset S11** GO enrichment of FTDEGs at 1 dpi (*Hpa* vs. mock) in Col-0 roots.

**Dataset S12** GO enrichment of FTDEGs at 2 dpi (*Hpa* vs. mock) in Col-0 roots.

**Dataset S13** GO enrichment of FTDEGs at 3 dpi (*Hpa* vs. mock) in Col-0 roots.

**Dataset S14** GO enrichment of FTDEGs at 4-5 dpi (*Hpa* vs. mock) in Col-0 roots.

**Dataset S15** GO enrichment of FTDEGs at 6-7 dpi (*Hpa* vs. mock) in Col-0 roots.

**Dataset S16** Transcription factor binding site analysis (SEA) on DREM clusters.

**Dataset S17** Transcription factor binding site analysis (SEA) on Splinecluster clusters

**Dataset S18** Transcription factor binding site analysis (DAP-seq) on DREM clusters.

**Dataset S19** Transcription factor binding site analysis (DAP-seq) on Splinecluster clusters.

**Dataset S20** Description of TFs from transcription factor binding site analysis.

**Dataset S21** Two-way ANOVA analysis of genotype and treatment effects on resistance against *Hpa* in responding Col-0 plants growing in soil conditioned by different mock- or *Hpa*-treated genotypes.

## Supplemental Figures and Tables Qi et al., 2026

**Fig. S1.**
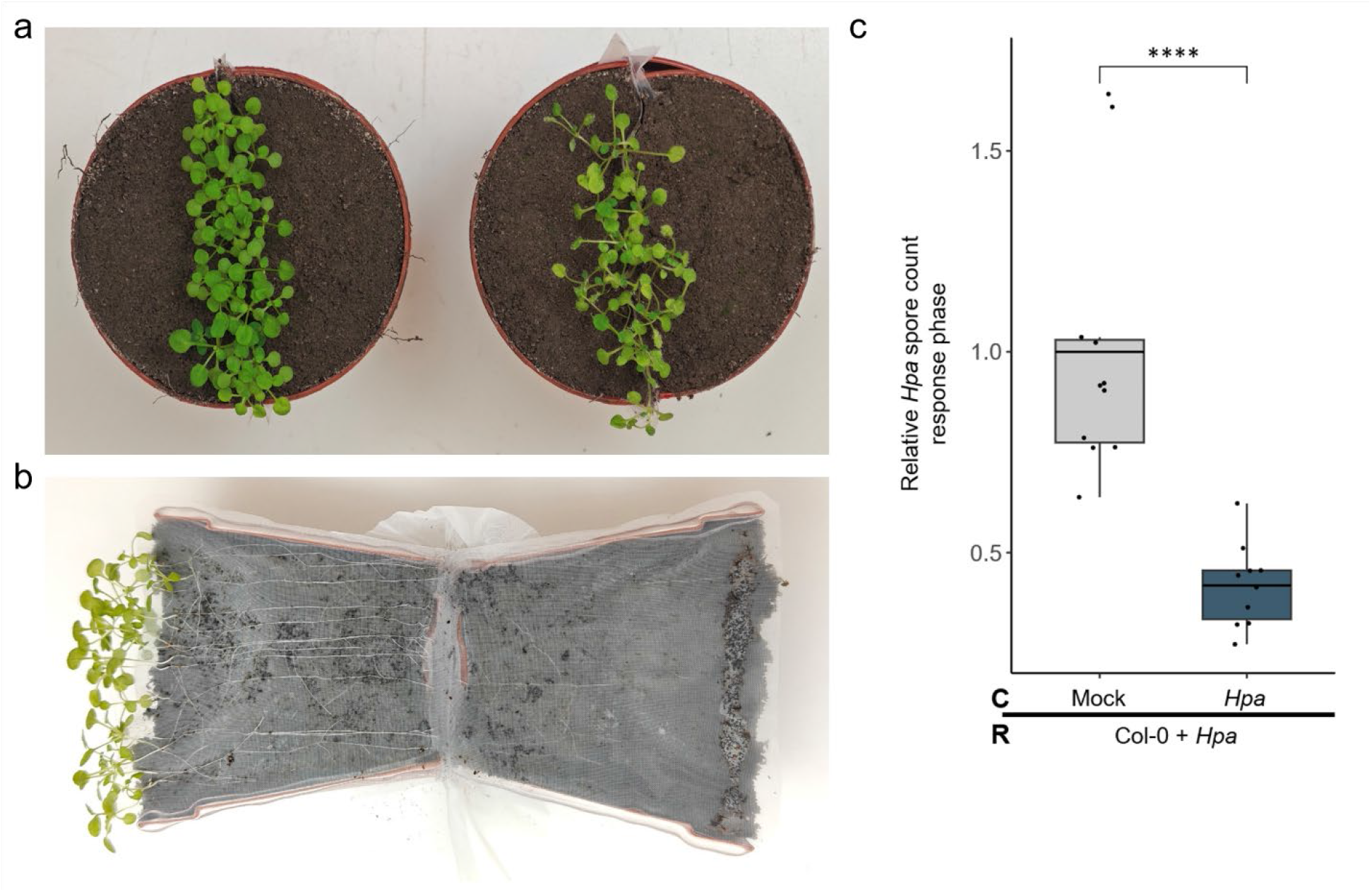
Tea bag growth system used for studying *Hpa*-induced SBL establishment. (a) Pots with 3-week-old Arabidopsis plants growing within in a nylon mesh pocket surrounded by Reijerscamp soil in the pot (the tea bag system). The left pot shows mock-treated conditioning plants and the right pot shows *Hpa*-infected conditioning plants. (b) Inside view of the tea bag system with 3-week-old seedlings showing intact, clean roots. (c) The “tea bag” system preserves the establishment of *Hpa*-induced SBL. Spore production of *Hpa* in response (R) populations of Col-0 plants growing in the mesh pocket of the “tea bag” system conditioned (C) by populations of wild-type Col-0 growing within the mesh pocket and that were mock- or *Hpa*-treated for 7 days (these conditioning pots are pictured in (a)). Relative *Hpa* spore counts represent the combined results of independent experiments, with spore counts normalized to the mock-treated conditioning group within each experiment. One-tailed Student’s t-test; **** *P* < 0.0001; *n* = 10-11 pots per experiment.

**Fig. S2.**
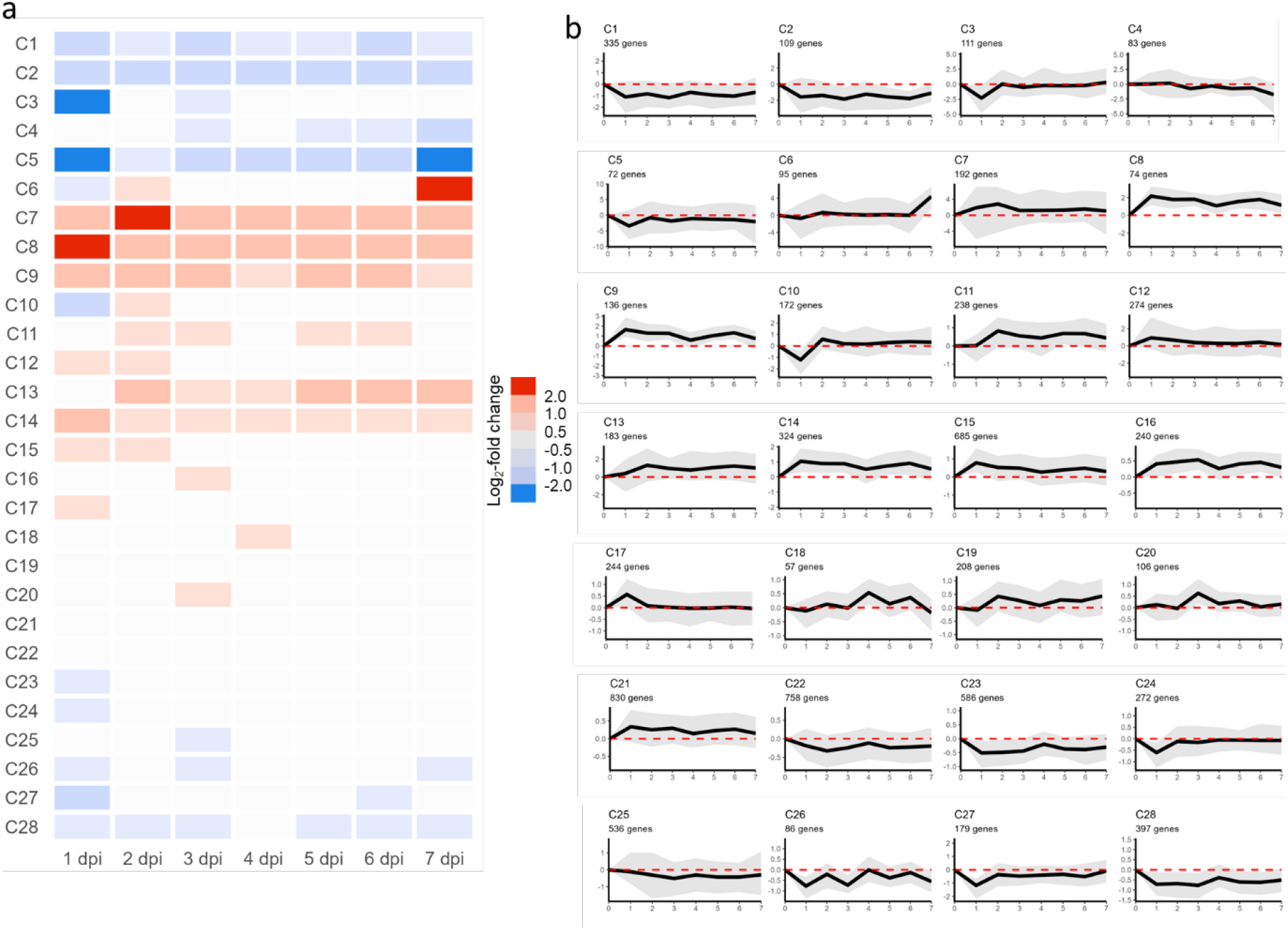
Clustering of temporally co-expressed DEGs in roots after foliar *Hpa* treatment. (a) The set of 7498 differentially expressed genes (DEGs) over a 7-day time course following *Hpa* inoculation was clustered based on their expression patterns (log_2_-fold change of *Hpa*/mock) using SplineCluster, resulting in 28 distinct clusters (C1-C28). Each row of the heatmap represents the average log_2_-fold change (*Hpa*/mock). (b) Individual cluster profiles over the 7-day time course. Shown is the mean log_2_-fold change of *Hpa*/mock of all genes in a cluster (black line), with gray error bars indicating the standard deviation. Cluster number and cluster size are denoted above each plot.

**Fig. S3.**
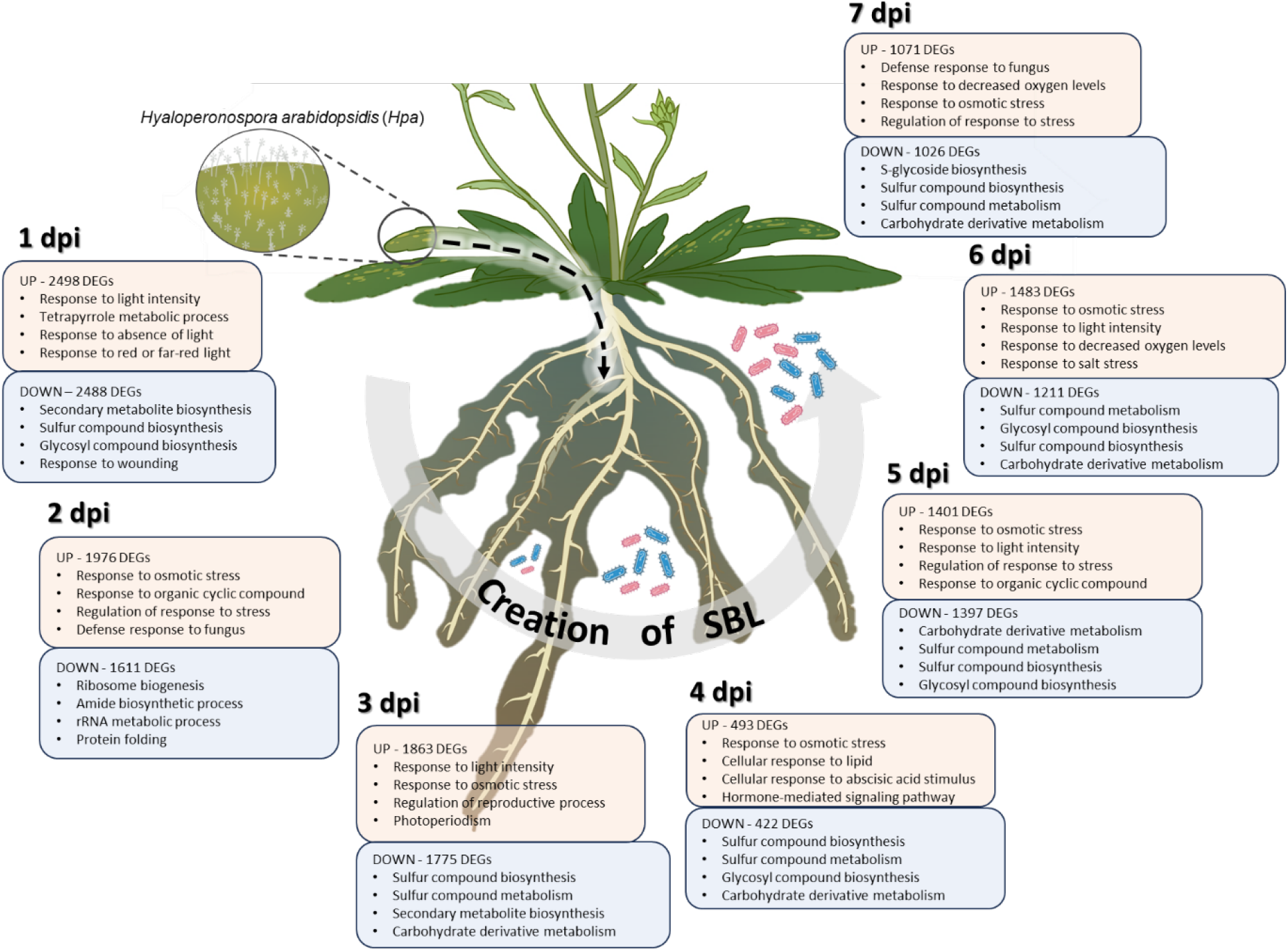
GO enrichment results of DEG analysis per time point in Arabidopsis roots upon infection by *Hpa*. Genes that are differentially expressed per time point were used for GO enrichment analysis. DEGs are categorized by upregulation (UP) and downregulation (DOWN). The most significantly enriched 4 biological process GO terms are shown for each category.

**Fig. S4.**
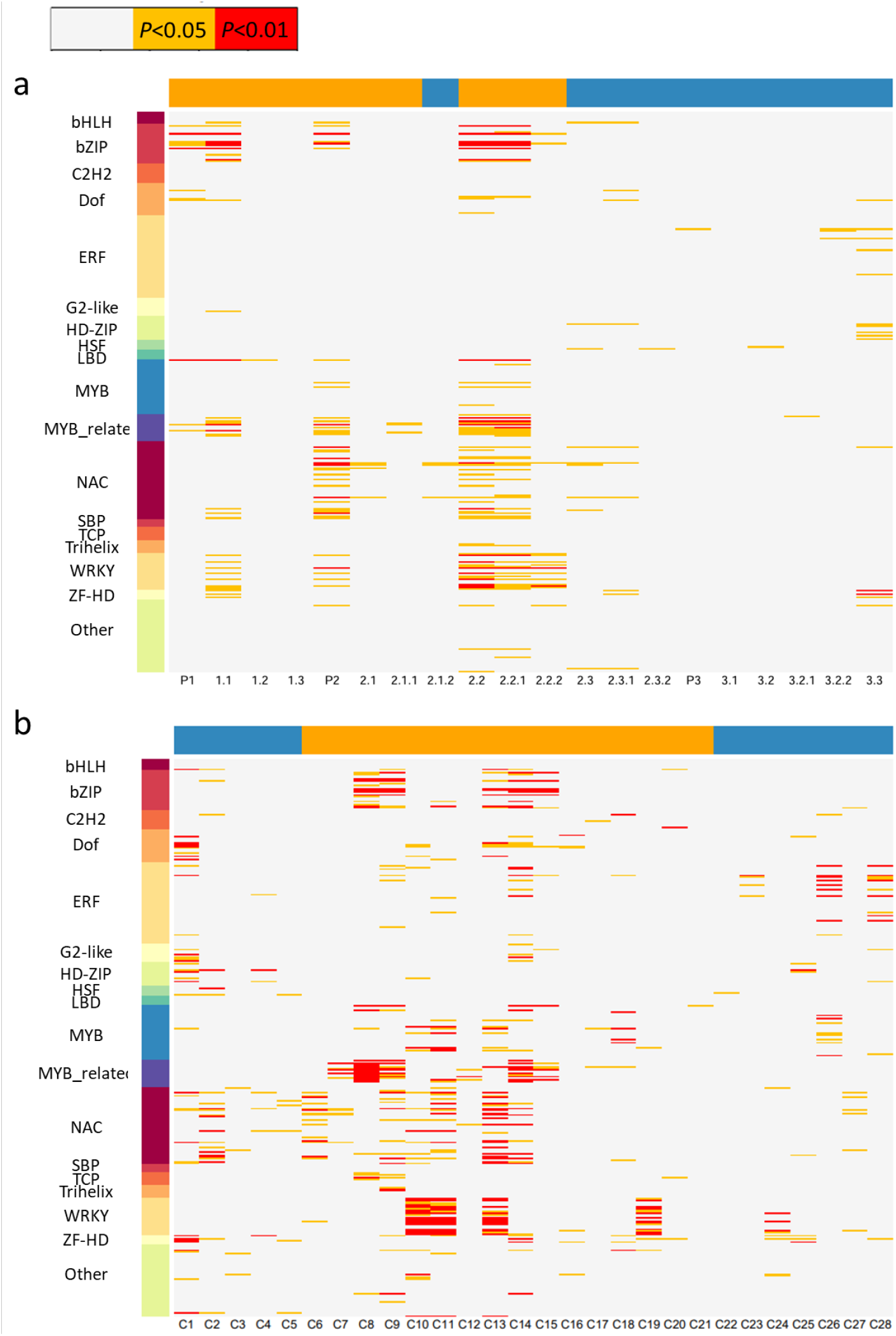
Enrichment of transcription factor binding sites in temporally co-expressed DEGs in roots after *Hpa* foliar treatment. (a) Enrichment of transcription factor binding sites in paths from the DREM analysis (see Fig. **2a**). (b) Enrichment of transcription factor binding sites in clusters from the SplineCluster analysis (see Fig. **S3**). Transcription factor binding sites were inferred from the filtered DAP-seq dataset. Colors denote levels of statistical significance based on a hypergeometric test (yellow means *P* < 0.05, red means *P* < 0.01). Colored bar above the heatmap indicates if genes in the cluster are on average upregulated (yellow) or downregulated (blue) after *Hpa* treatment. Only the significant transcription factor binding sites are depicted. TFs within each family were ordered in the figure based on presence/absence of binding sites in the clusters.

**Fig. S5.**
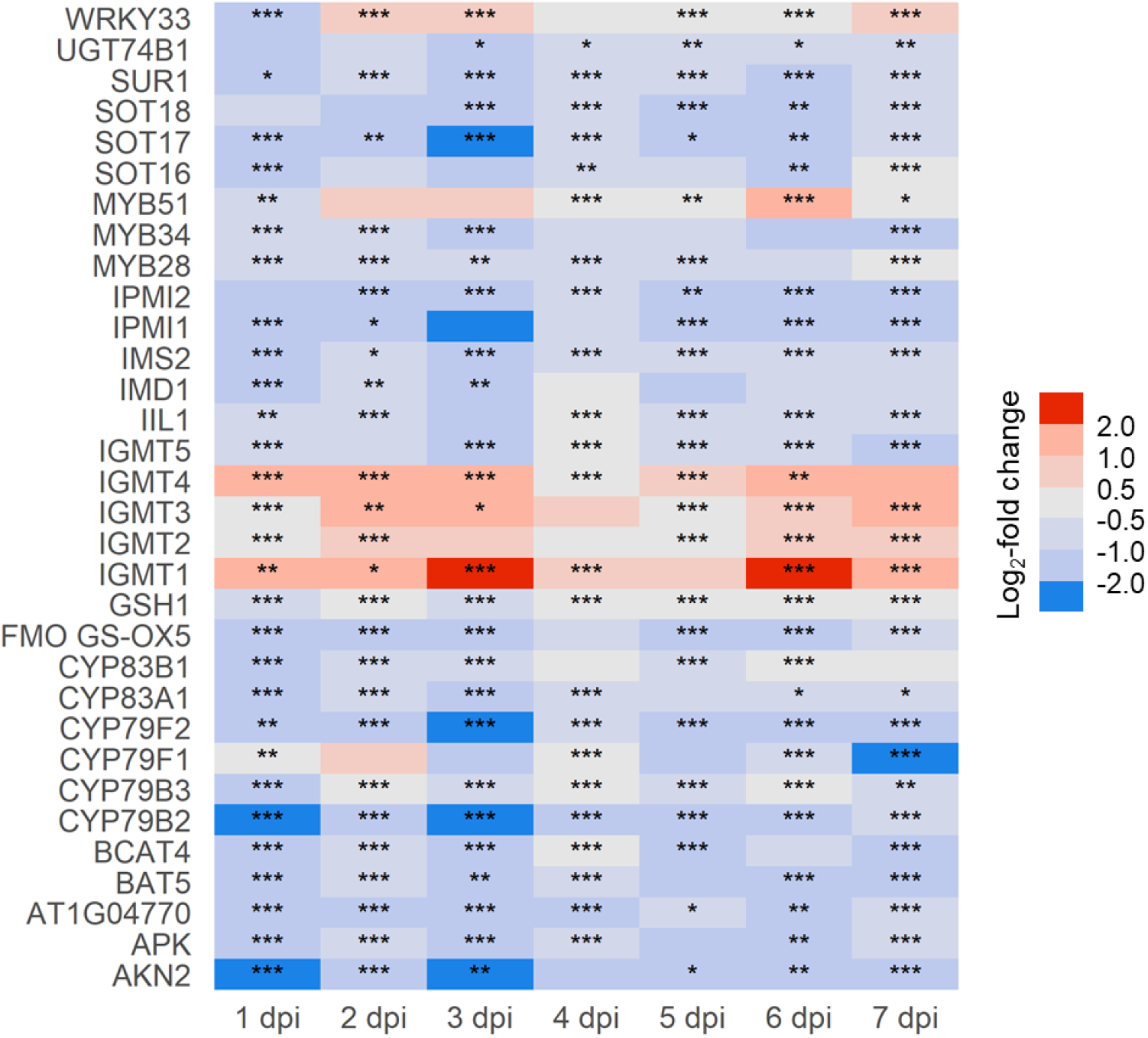
Expression profile of DEGs involved in glucosinolates biosynthesis process. Temporal expression profiles of genes annotated with GO term “glucosinolates biosynthetic process” following *Hpa* inoculation, including both IGS and AGS biosynthesis genes. Rows indicate the log_2_-fold change between *Hpa* and mock treatments at 1-7 dpi, following the color range indicated in the legend. Asterisks indicate significant differential expression based on DESeq2 analysis (* *P* < 0.05, ** *P* < 0.01, *** *P* < 0.001).

**Fig. S6.**
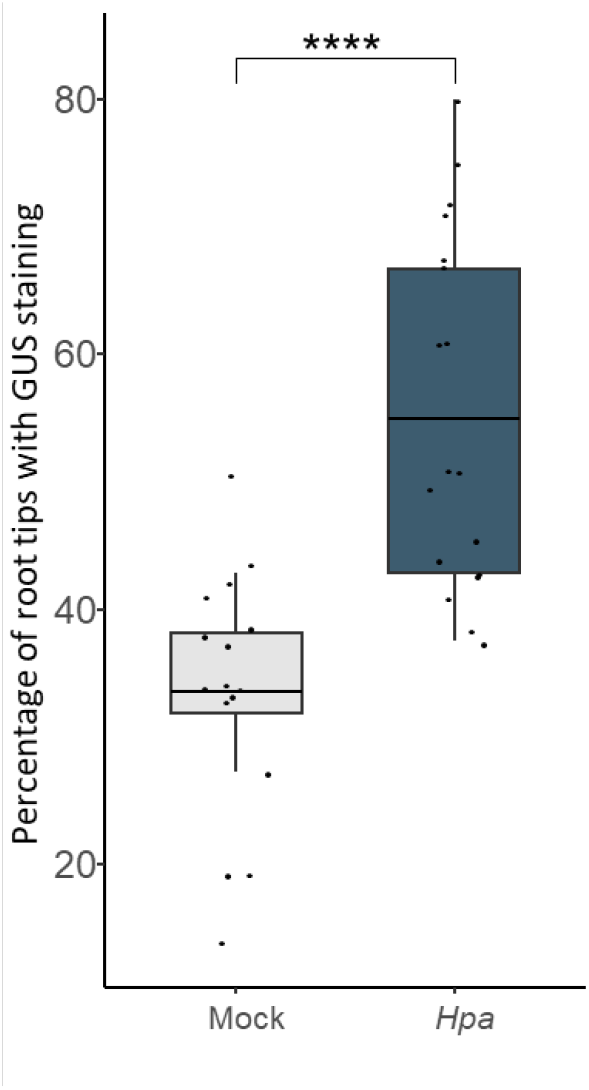
Quantification of the percentage of root tips showing GUS staining per plant of *WRKY33pro:GUS* reporter line under mock and *Hpa*-treated conditions, grown *in vivo* in the “tea bag system”, at 3 dpi. Asterisks indicate statistically significant differences (Student’s t-test; **** *P* < 0.0001; *n* = 16). For representative pictures of the GUS staining, see Fig. 5b.

**Fig. S7.**
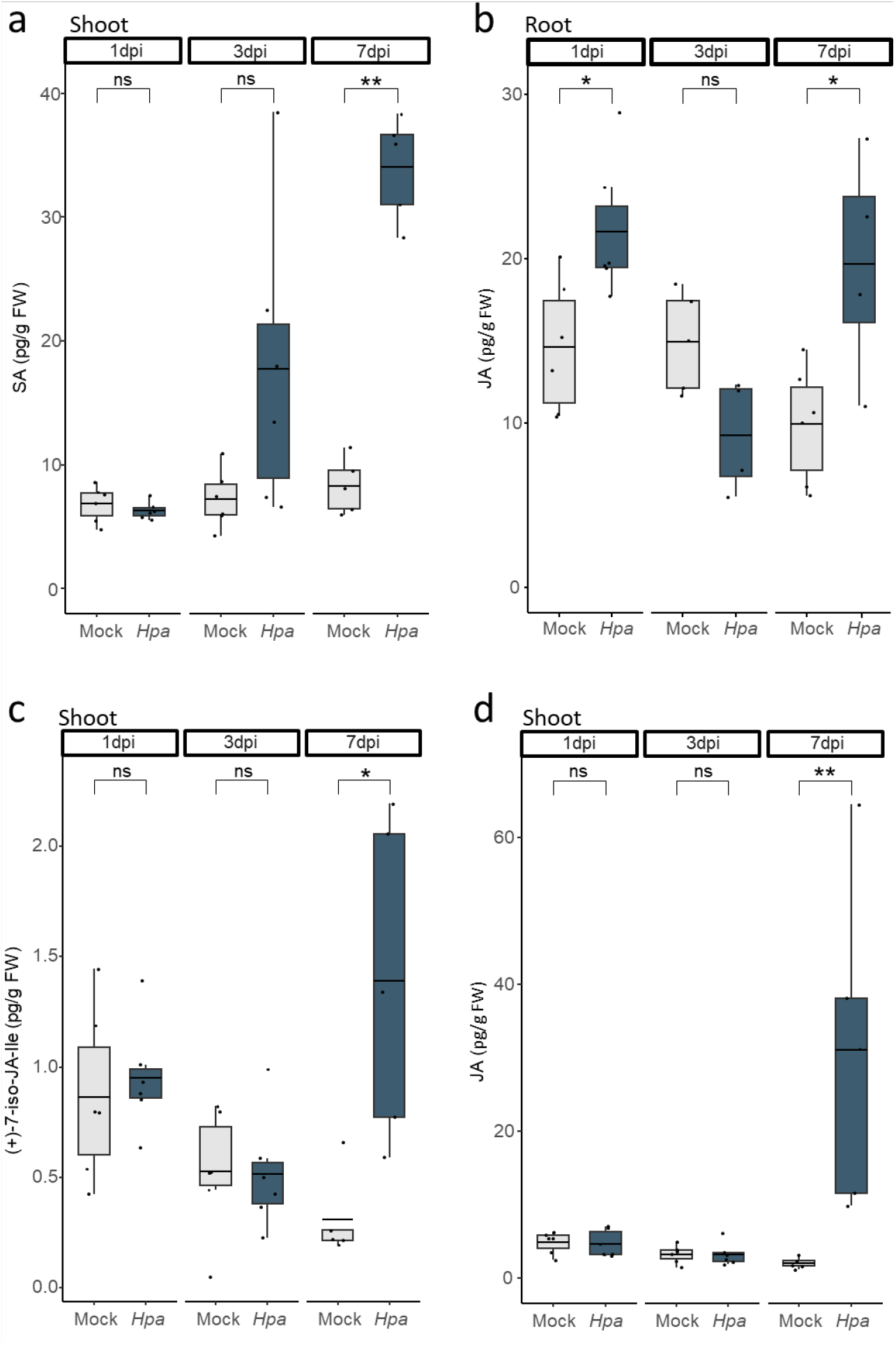
Measurement of SA and JA in Arabidopsis shoots and roots. (a) Concentration of the hormone SA in shoots at 1, 3 and 7 dpi after foliar mock or *Hpa* treatment. (b) Concentration of the hormone JA in roots at 1, 3 and 7 dpi after foliar mock or *Hpa* treatment. (c) and (d) Concentrations of the hormones (+)-7-iso-JA-Ile and JA in shoots at 1, 3 and 7 dpi after foliar mock or *Hpa* treatment. Hormone levels are normalized to fresh weight (FW) of each sample. Asterisks indicate statistically significant differences (Student’s t-test; * *P* < 0.05, ** *P* < 0.01; *n* = 6).

**Fig. S8.**
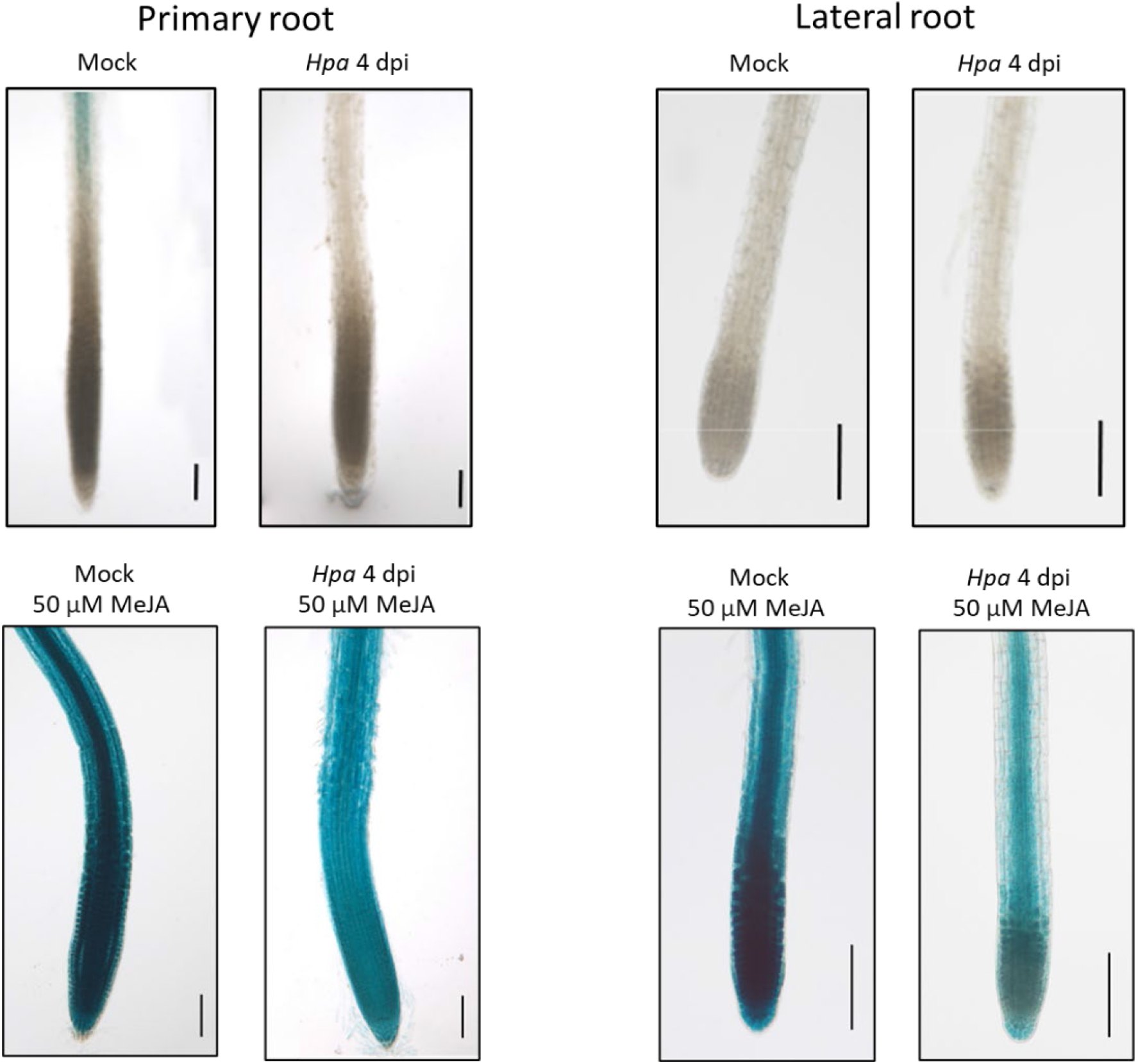
GUS staining of *in vitro* grown *JAZ10pro:GUS* reporter line in root tips of *Hpa*- and MeJA-treated plants. Figures show representative images of primary root tips (left) and lateral root tips (right) under the following conditions: Leaf mock and *Hpa* treatment at 4 dpi (upper panel), leaf mock or *Hpa* treatment at 4 dpi + root 50 μM MeJA treatment for 3 h (lower panel). The pictures show representative root tips grown *in vitro.* Scale bars = 100 μm.

**Table S1.** Arabidopsis lines used in this study.

| Line name | Mutant allele / Transgene | Genetic Background | Stock ID | Source | Reference |
| --- | --- | --- | --- | --- | --- |
| Col-0 | Wild type | Columbia (Col-0) | — | Laboratory stock | — |
| *wrky33-1* | T-DNA insertion mutant | Col-0 | SALK_006603 | NASC | Zheng et al., 2006 |
| *cyp81f2* | T-DNA insertion mutant | Col-0 | SALK_073776 | NASC | Pfalz et al., 2009 |
| *cyp81f4* | T-DNA insertion mutant | Col-0 | SALK_024438 | NASC | Pfalz et al., 2011 |
| *tt3-5* | Mutant allele | Col-0 | — | NASC | Peer et al., 2001 |
| *tt4-3* | Mutant allele | Col-0 | — | NASC | Peer et al., 2001 |
| *pad3-1* | Mutant allele | Col-0 | — | Laboratory stock | Zhou et al., 1999 |
| *myb28-1 myb29-1* | T-DNA insertion double mutant | Col-0 | — | Laboratory stock | Sønderby et al., 2007 |
| *npr1-1* | Mutant allele | Col-0 | — | Laboratory stock | Cao et al., 1997 |
| *sid2-1* | Mutant allele | Col-0 | — | Laboratory stock | Wildermuth et al., 2001 |
| *myc2 (jin1-7)* | Mutant allele | Col-0 | — | Laboratory stock | Berger et al., 1996 |
| WRKY33pro:GUS | GUS reporter line | Col-0 | — | Provided by Paul Schulze-Lefert | Lippok et al., 2007 |
| JAZ10pro:GUS | GUS reporter line | Col-0 | — | Provided by Edward Farmer | Acosta et al., 2013 |

**Table S2.**
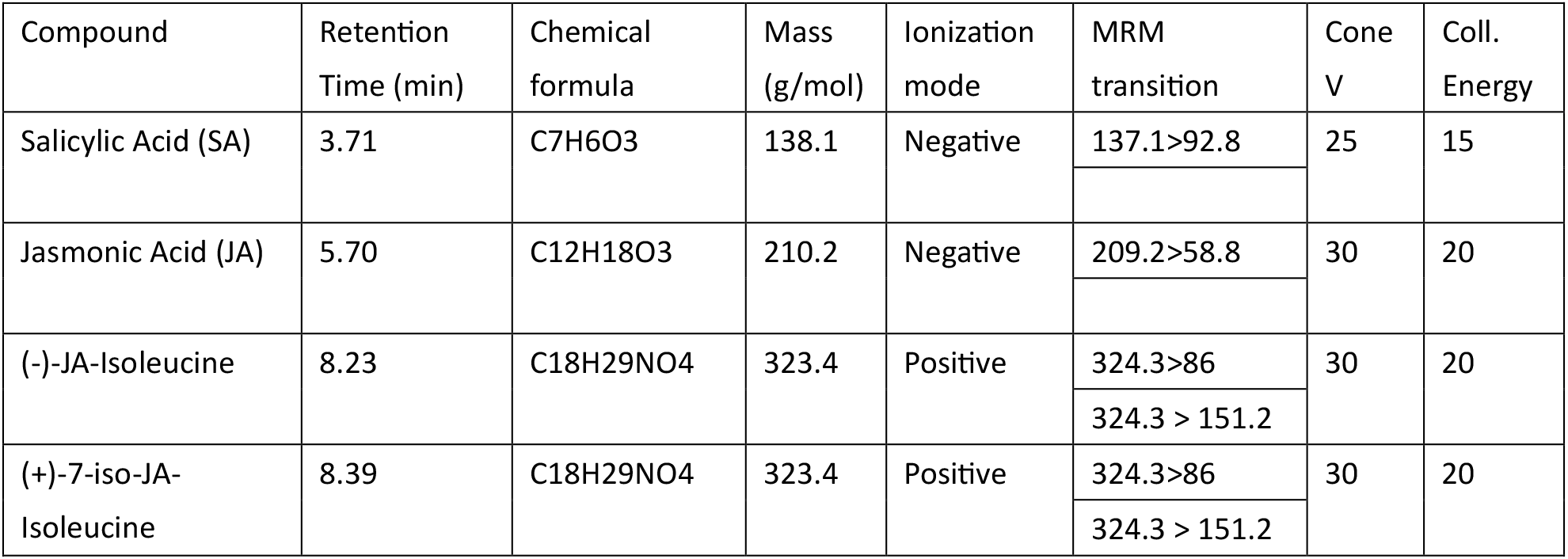
Multiple reactions monitoring (MRM) transitions table. Transitions for all hormones and corresponding internal standards used in this study.

| Compound | Retention Time (min) | Chemical formula | Mass (g/mol) | Ionization mode | MRM transition | Cone V | Coll. Energy |
| --- | --- | --- | --- | --- | --- | --- | --- |
| Salicylic Acid (SA) | 3.71 | C7H6O3 | 138.1 | Negative | 137.1>92.8 | 25 | 15 |
| Jasmonic Acid (JA) | 5.70 | C12H18O3 | 210.2 | Negative | 209.2>58.8 | 30 | 20 |
| (-)-JA-Isoleucine | 8.23 | C18H29NO4 | 323.4 | Positive | 324.3>86 | 30 | 20 |
|  |  |  |  |  | 324.3 > 151.2 |  |  |
| (+) -7-iso-JA-Isoleucine | 8.39 | C18H29NO4 | 323.4 | Positive | 324.3>86 | 30 | 20 |
|  |  |  |  |  | 324.3 > 151.2 |  |  |

